# Rapid automated 3-D pose estimation of larval zebrafish using a physical model-trained neural network

**DOI:** 10.1101/2023.01.06.522821

**Authors:** Aniket Ravan, Ruopei Feng, Martin Gruebele, Yann R. Chemla

## Abstract

Quantitative ethology requires an accurate estimation of an organism’s postural dynamics in three dimensions plus time. Technological progress over the last decade has made animal pose estimation in challenging scenarios possible with unprecedented detail. Here, we present (i) a fast automated method to record and track the pose of individual larval zebrafish in a 3-D environment, applicable when accurate human labeling is not possible; (ii) a rich annotated dataset of 3-D larval poses for ethologists and the general zebrafish and machine learning community; and (iii) a technique to generate realistic, annotated larval images in novel behavioral contexts. Using a three-camera system calibrated with refraction correction, we record diverse larval swims under free swimming conditions and in response to acoustic and optical stimuli. We then employ a convolutional neural network to estimate 3-D larval poses from video images. The network is trained against a set of synthetic larval images rendered using a 3-D physical model of larvae. This 3-D model samples from a distribution of realistic larval poses that we estimate a priori using a template-based pose estimation of a small number of swim bouts. Our network model, trained without any human annotation, performs larval pose estimation with much higher speed and comparable accuracy to the template-based approach, capturing detailed kinematics of 3-D larval swims.

**Author Summary:** Larval zebrafish swimming has been studied extensively in 2-D environments, which are restrictive compared to natural 3-D habitats. To enable rapid capture of 3-D poses, we collect three orthogonal video projections of swim behaviors in several behavioral settings and fit poses to a physical model. We then use the physical model to generate an auto-annotated stream of synthetic poses to train a convolutional neural network. The network model performs highly accurate pose predictions on over 600 real swim bouts much faster than a physical model fit. Our results show that larvae frequently exhibit motions inaccessible in a 2-D setup. The annotated dataset could be used by ethologists studying larval swimming dynamics, and by the machine learning community interested in multi-dimensional time series and 3-D reconstruction. Using the ability to render images with multiple synthetic poses, our method can be extended to collective behavior.

## Introduction

Neuroethologists have long sought an understanding of the neural basis of behavior, through multiple model organisms offering varying degrees of complexity of their nervous system. An important step in achieving this goal is to develop quantitative tools to objectively identify distinct behaviors (1–8). This step requires accurate tracking of an organism’s pose as a function of time. Historically, this has been done by manually labeling the points of interest on the body, which is a laborious process and subject to errors. However, over the last decade, it has become increasingly possible to automate several image analyses tasks, including animal pose estimation with human-level accuracy (9–19). These advances can be credited to the success of artificial neural networks in conjunction with development graphical processing units (GPUs). Artificial neural network models can be tuned for pose estimation, given examples of image data with a priori annotated poses. Since these annotations are created by manual human labelling in most cases, the network’s accuracy is bound by that of the source generating the annotations.

*Danio rerio* (zebrafish) larvae are a popular choice of model vertebrate organism and their swimming behavior in response to several environmental stimuli has been studied extensively over the last several decades (5,6,20–29). These studies are mostly restricted to 2-D, where the larva is constrained to swim in a shallow (2 – 3 mm height) medium and imaged using a single overhead camera. Thus, these measurements lack a complete three-dimensional picture of larval swimming dynamics. Moreover, the depth available for swimming is not only smaller than the length of the organism (3 – 4 mm long at 6-7 days post fertilization (dpf)), but also severely restrictive compared to its native environment (30). It is plausible that this constraint influences the swimming motion of the larva. Recent work (31) showing differences in neural activity of a constrained vs. a freely moving *C. elegans* makes a strong case for studying unconstrained animals.

These limitations motivated us to develop a fast and accurate technique for estimation of the poses (3-D position, orientation, and shape of the backbone) of larval zebrafish swimming in a 3-D environment. We collected videos of individual larvae swimming in a cubic glass tank of spatial dimensions an order of magnitude larger than the length scale of a larva. Videos were captured by three orthogonal cameras, calibrated to account for non-linearities due to refraction (**Materials and Methods(a) Instrumentation:** *Camera calibration*). Our fully automated method of pose estimation relies on a convolutional network trained on a set of digitally rendered larval images with annotated pose (**Figure 1**). We refer to these rendered images as “physical model images” henceforth.

**Figure 1:**
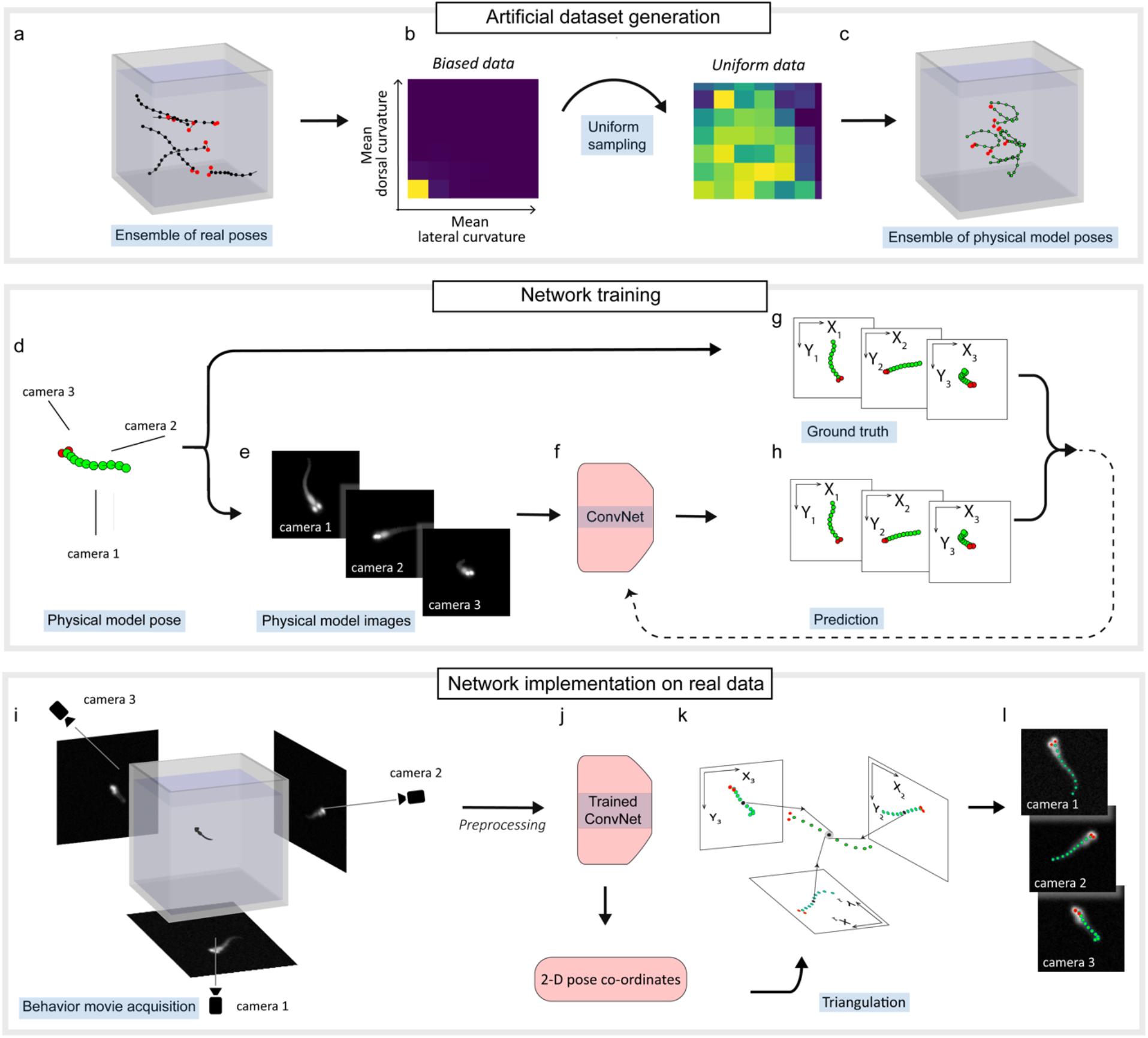
Convolutional neural network model trained on physical model images performs fast and accurate pose estimation on real images. (a) Ensemble of real poses: An ensemble of real poses (N=35714) is generated by estimating larval poses using a template-based pattern recognition approach and selecting a subset of poses estimated with high accuracy. (b) *Biased data:* The ensemble of real poses is biased towards larval shapes with nearly straight backbone. This is visualized on a coarse grained 2-D space of mean lateral and dorsal body curvature. The panel shows a histogram of mean lateral curvature and mean dorsal curvature computed over the ensemble of real postures. A majority of the poses are seen to lie close to (0,0), which corresponds to larvae with a nearly straight backbone. *Uniform data:* A subset of 2500 poses is sampled from the ensemble of real poses, such that larval shapes with different lateral and dorsal curvature are represented uniformly. Samples from the ensemble of real pose are chosen conditional on a probability distribution function inversely proportional to the probability density of the biased data (b-*Biased data*). (c) Ensemble of physical model poses: An ensemble of physical model poses (*N* = 500,000) drawn using a probabilistic model generated from the uniform data (*c-Uniform data*) of 2500 real poses is generated. Each physical model pose is drawn from a probability distribution of backbone angles, roll and inclination, appending the position and yaw chosen uniformly at random. (c) Physical model pose: Every physical model pose in the ensemble of physical model poses in (c) is used to generate the training dataset to train the neural network model. (e) Rendering physical model images for input to the network model: A physical model of the larva is used to render physical model images of size 141×141 corresponding to the sampled pose in (d). (f) Convolutional neural network model: Physical model images from (e) are passed as three input channels to the convolutional neural network. (g) Generation of ground truth annotations: 2-D pose annotations are generated by mapping the 3-D pose using a nonlinear projection function estimated during camera calibration. (h) Neural network training: The neural network predictions are designed to be 3×12 2-D pose coordinates. The network parameters are tuned using the loss function defined as the root mean squared error between the network predictions and the ground truth annotations. (i) Larval images recorded by three orthogonal cameras are preprocessed by cropping and background subtraction. (j) The preprocessed images are passed to the trained neural network to obtain 2-D pose coordinates of the recorded larva. (k) Triangulation: 3-D pose coordinates are triangulated using the 2-D pose coordinates and the empirically determined projection function. (l) The 2-D projections of the estimated pose are superimposed on a set of real images for visual inspection.

The physical model images are rendered using a voxel-based physical model of the larva, where voxels are mapped to pixels on three orthogonal camera views using 3-D-to-2-D projection parameters obtained during camera calibration. The poses used to generate these images are sampled from a distribution of poses estimated a priori using a computationally expensive template-based pose estimation (**Materials and Methods(b) Template-based pose estimation**). The use of annotated physical model images obviates the need for human labelling, which in this context is not merely an inconvenience, but also extremely difficult. The challenges in human labelling are due to occlusions and the necessity to label consistent points on the organism across the three views and across different frames on the larval backbone. Such consistency in labelling is made challenging by the lack of any natural fiducial markers on the organism. A similar approach involving digitally generated annotated images via the use of real images and affine transforms was recently developed (12) for pose estimation of *C*. *elegans*. However, this approach cannot be extended to the case of a 3-D larva where larval images are frequently characterized by self-occlusions. Moreover, zebrafish larvae consist of more complex features than worms, like eyes, head and belly, whose motion cannot be accounted for by using affine transforms of their 2-D projections.

Here, we record larval images in three commonly studied behavioral contexts: free swimming, acoustic startle, and dark flash (**SI Figure 2**) and use our physical model-trained neural network tool to estimate larval poses in these recordings. We then evaluate the convolutional neural network’s performance using the correlation coefficient between the physical model images resulting from the model’s predictions and raw images. The neural network model produces accurate results much faster than the template-based pattern recognition technique. Our measurements reveal a rich set of 3-D kinematics of larval swims not accessible in classical 2-D experiments.

Our work can be summarized by three modular tools presented here as a single technique for larval pose estimation:

1. A neural network model for fast and accurate 3-D pose estimation of zebrafish larvae
2. A large collection of diverse larval 3-D poses and the corresponding raw images, of interest to the ethologists, zebrafish researchers, and machine learning community
3. A technique to render realistic larval images which can be used to generate an arbitrarily large training dataset for a neural network in novel behavioral contexts like collective behavior

A visualization of a 3-D reconstructed larval swim bout from an acoustic startle experiment is presented in **Supplementary Video 1**.

## Results

### Convolutional neural network model trained on physical model images performs fast and accurate pose estimation on real images

#### Experimental design

Fish swimming measurements were carried out on 6-9 days-old larvae a few millimeters in length in a cubic glass tank of dimensions 7 x 7 x 7 cm (see **SI Figure 2a** and **Materials and Methods(a) Instrumentation**). Movies of the zebrafish were obtained using 3 synchronized high-speed cameras taking orthogonal images of the fish tank at 500 frames per second (fps) (**SI Figure 1a**). While two orthogonal axes are in principle sufficient to reconstruct the position of simple rod-like shapes, three axes provide redundancy to correct for small errors due to water refraction, and additional information for more complex shapes, such as a highly bent backbone that obscures parts of itself from certain view angles. We calibrated the cameras taking into account refraction at the water-glass and glass-air interfaces using a dot grid. We also estimated a projection function, modelled as a cubic polynomial, that maps 3-D coordinates in the tank to 2-D pixel coordinates for each camera (see **Materials and Methods(a) Instrumentation:** *Camera calibration: Mapping from lab to camera coordinates* and **SI Figure 3**). A 3 x 3 x 3 cm cubic region in the center of the tank overlapped the field of view of all the three cameras. For all our experiments, we recorded larvae swimming in this cubic region. We used three orthogonally placed and diffused high-power (50 W) near-infrared LEDs (850 nm) for illumination outside the larvae’s wavelength sensitivity (32). This illumination did not heat the water significantly yet compensated for the high frame rate necessary to capture maneuvers reliably and the small optical aperture (large *f*-ratio) necessary in order to achieve a large depth of field.

Zebrafish larvae were placed in the tank and allowed to swim in free swimming, acoustic startle, and dark flash contexts. We collected a total of 630 movies of larvae swimming in 3-D, with at least 100 movies from each type of experiment (free swimming – 303, acoustic startle – 162, dark flash – 165). For acoustic startle experiments, an acoustic stimulus was generated by dropping a weight from a fixed height onto the platform mounting the glass tank (**SI Figure 2c**). A dark flash stimulus was generated by turning off a white LED placed near the tank for 10 s (**SI Figure 2b**). To increase measurement throughput, both types of stimuli were triggered automatically whenever the three cameras detected at least one larva in the 3 x 3 x 3 cm imaging region. Consecutive stimuli were separated by at least 4 minutes in time to prevent habituation of the larvae to the stimulus. This inter-stimulus interval (ISI) is significantly larger than an ISI of 15 seconds reported in (33) to induce habituation to acoustic startle stimuli. We found in our experiments that an ISI of 4 minutes was sufficient to obtain consistently larval responses to dark flashes in the form of O-bends (Burgess & Granato, 2007c). The setup is described in detail in **Materials and Methods(a) Instrumentation** and **SI Figure 2**.

#### The 3-D physical model of a larval zebrafish

We represented the larva using a physical model. The physical model requires 12 3-D coordinates to define a larval pose: 10 equally spaced coordinates along its backbone, and 2 coordinates defining the centroid of its eyes. The physical model is composed of 9 segments connected by flexible hinges, and includes a larval anterior composed of head, eyes, and ventral region located by the first two segments and a posterior composed of seven tail segments (**SI Figure 1c**). The coordinates of the physical model are then used to render realistic 2-D larval projections (see **Materials and Methods(b) Template-based pose estimation**: *Rendering larval projections, Physical model of the larva* for more details). We refer to these projections as “physical model images”.

A definition of the physical model in terms of the angles between its consecutive segments is more suitable for direct quantitative analysis of behavior. Thus, we also formulated an equivalent description of the physical model in 3-D by specifying a set of adjustable parameters encoded in the 22-parameter vector **p** = (*x*_0_, *y*_0_, *z*_0_, *θ*_0_, Δ*θ*_1_,… Δ*θ*_8_, *φ*_0_, Δ*φ*_1_,… Δ*φ*_8_, *γ*_0_) and a fixed parameter, *l* (fish length). *x*_0_, *y*_0_, *z*_0_ determine the centroid of the head relative to the lab reference frame *x*_lab_, *y*_lab_, *z*_lab_, and the Euler angles *θ*_0_, *φ*_0_, *γ*_0_ determine the head orientation (yaw, inclination, and roll, respectively) in the lab reference frame. **θ** = (Δ*θ*_1_,… Δ*θ*_8_, Δ*φ*_1_,… Δ*φ*_8_) is the set of angles that give the orientation of each body segment and determines the fish shape. Here, Δ*θ_i_* and Δ*φ_i_* are the bending angles for the (*i*+1)^th^ segment in the lateral and dorsal planes, respectively, as measured in the fish reference frame *X, Y, Z*, defined by the head orientation *θ*_0_, *φ*_0_, *γ*_0_ (see **SI Figure 1c** for angle nomenclature).

A larval pose can thus be uniquely identified using either of the following equivalent representations:

1. 3-D pose coordinates: 12 3-D coordinates in the cartesian lab reference frame *x*_lab_, *y*_lab_, *z*_lab_ that define the physical model
2. 2-D projection pose coordinates: 2-D projections of the 3-D pose coordinates obtained using the projection function
3. Parameter vector **p** and *l*

#### Generation of the training and validation datasets

We sought to train a convolutional neural network model to estimate larval poses from pre-processed larval images (see **Materials and Methods(c) Neural Network pose estimation**: *Preprocessing training dataset*) captured by the three cameras. Our approach is summarized in **Figure 1**, and consists of three steps: generation of a synthetic training dataset, training of the convolutional neural network, and implementation of the network on real larval images. In the last step, the network maps the larval images to three 2-D projection pose coordinates, which can then be triangulated to obtain the 3-D pose coordinates using the projection function in a single post-processing step.

**Figure 1a-c** illustrates the procedure for generating training and validation datasets. These datasets consist of physical model images used as the neural network inputs and their corresponding 2-D projection pose coordinates as the ground truth annotations. We rendered 500,000 physical model images and computed their corresponding 2-D projection pose coordinates using an ensemble of physical model poses (see *Ensemble of physical model poses* in **Figure 1c**), which is a set of realistic larval poses generated using a probabilistic model informed from an ensemble of real larval poses (see **Figure 1a-b**). We split the training and validation dataset in a 9:1 ratio while training the convolutional neural network model. The lack of human intervention in generating this training and validation dataset results in efficient generation of accurate labels.

The ensemble of real larval poses (**Figure 1a**) required to generate the training dataset comprises 35714 distinct real larval poses. It is obtained using a computationally expensive template-based pose estimation performed in over 424 larval video recordings or 41756 frames (see **SI Figure 1 and Materials and Methods(b) Template-based pose estimation)** and selecting a subset of the poses with a pose estimation score of at least 0.9 (see **Results: *Model evaluation*** for a definition of pose estimation score). Briefly, we perform a search in the 22-parameter space of the vector **p**. For any given **p** and *l*, the physical model is uniquely determined, and physical model images are rendered using the projection function (see **Materials and Methods(b) Template-based pose estimation:** *Physical model of the larva*). We optimize **p (see Materials and Methods(b) Template-based pose estimation:** *Optimization*) by minimizing the sum of squared difference between the physical model images and the pre-processed data captured by the orthogonal cameras (**SI Figure 1b-h)**.

Finally, the ensemble of real larval poses is used to generate a realistic and unbiased ensemble of physical model poses using a probabilistic model (**Figure 1a-c**). This approach is adopted mainly for two reasons: (1) We observe that a large fraction of real larval poses consist of a nearly straight backbone poses, and this biased ensemble is thus not ideal as a training dataset. The distribution of *mean lateral curvature* <|Δ*θ*_i_|> and *mean dorsal curvature* <|Δ*φ*_i_|>, where *i ϵ* [1,8] (**Figure 1b**-*Biased data*) illustrates this point, showing that most swim bouts are distributed near <|Δ*θ*_i_|> = 0 and <|Δ*φ*_i_|> = 0). (2) A probabilistic model informed from the ensemble of real poses allows us to generate an arbitrarily large number of distinct physical model poses. This model is obtained in two steps. First, we use a kernel density estimate to create a uniform data set of 2500 larval poses, sampled from the entire set of 35714 poses in the ensemble of real larval poses, such that larval shapes of different lateral curvature <|Δ*θ*_i_|> and dorsal curvature <|Δ*φ*_i_|> are uniformly represented (depicted in **Figure 1b** as the transformation from *Biased data* to *Uniform data* and described in **Materials and Methods: Neural network pose estimation:** *Generation of ensemble of physical model poses*). The size of the uniform data (*N* = 2500) was chosen as a trade-off between the diversity of larval poses and computational complexity. Next, we again use a kernel density estimate to generate the probability distribution 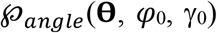 of all the 2500 poses in the uniform data. A physical model pose can be generated using a probabilistic model, which samples a vector (**θ**, *φ*_0_, *γ*_0_) from 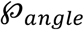 and assigns to it (*x*_0_, *y*_0_, *z*_0_) within the glass tank’s imaging volume and *θ*_0_ in the range (−*π, π*] by sampling from a uniform distribution. This probabilistic model is used to sample 500,000 novel and distinct larval poses, forming the ensemble of physical model poses (**Figure 1c**).

#### Training an artificial neural network model

The procedure for training the convolutional neural network is illustrated in **Figure 1d-h**. First, for each physical model pose in the ensemble (**Figure 1d**), physical model images are rendered using the voxel-based model (**Figure 1e** and **Materials and Methods(b) Template-based pose estimation**: *Physical model of the larva, Rendering larval projections*). These images have a modest resolution of 141×141, to accommodate completely larval projections of all sizes seen in our data. The larval projections are then displaced from the center of the image at random by 0-20 pixels and Gaussian white noise is added to mimic the experimental data (see **Materials and Methods(c) Neural network pose estimation:** *Preprocessing training dataset* for more details). At the same time, the 2-D projection pose coordinates are computed using the projection function and the 3-D pose coordinates, forming the ground truth annotations for the three images (**Figure 1g**).

We use a simple convolutional neural network (**Figure 1f**) inspired by the recent success of Residual Networks (34) on various pattern recognition tasks, including animal pose estimation (12,16). We leverage the highly effective feature-extraction power of residual blocks for our application. Our network consists of two modules: an encoder and a decoder. The encoder consists of four bottleneck residual blocks as implemented in (34) with 32, 64, 128, and 256 channels respectively. The decoder consists of three fully connected layers of dimensions 1×288, 1×144 and 1×72. The convolutional neural network model is described in greater detail in **Materials and Methods(c) Neural network pose estimation:** *Convolutional neural network model*.

The neural network accepts the three physical model images as three input channels (**Figure 1e,f**). The network’s prediction is designed to be 3×12 2-D coordinates, forming the predicted 2-D projection pose coordinates (**Figure 1h**). The network’s parameters are tuned by minimizing a loss function defined as the root-mean-squared-error between the predicted 2-D projection pose coordinates and the ground truth annotations. The loss function is made symmetric with respect to the two eyes (see **Materials and Methods(c)** *Neural network pose estimation* for more details on the loss function). Since the network model processes the three channels simultaneously, information between the three larval views is coupled for an accurate prediction of the 2-D projection pose coordinates. Thus, the lack of knowledge about the location of the larval backbone due to self-occlusion in one view can be compensated by other view(s) in which there is no self-occlusion to generate an accurate predicted annotation.

#### Pose inference using the artificial neural network model

The procedure for performing pose estimation using the trained convolutional neural network is illustrated in **Figure 1i-l**. The larval pose for each frame is performed independently. The raw data is first pre-processed by performing background subtraction (see **Materials and Methods(b) Template-based pose estimation**: *Preprocessing*) and cropping a window of 141×141 pixels around the three images of a larva, followed by median filtering (images shown in **Figure 1i**). These pre-processed images are passed to the network (**Figure 1j**) trained on physical model images to generate the 2-D projection pose coordinates (**Figure 1l**). The 12 points of the larval pose are triangulated in parallel using the inferred 2-D projection pose coordinates and the projection function (**Figure 1k**). The triangulation is trivial for the backbone coordinates. The eye coordinates are triangulated by taking into account the symmetry of the loss function with respect to the eye indices (see **Materials and Methods(c) Neural network pose estimation**: *Triangulation of 3-D pose coordinates*). We further improve the pose prediction by fitting a spline through the estimated 3-D backbone coordinates and computing 10 equally spaced coordinates along the arc length (35). This step ensures that the points estimated along the backbone are equidistant and the angles between the segments of the physical model (Δ*θ*_i_, Δ*φ*_i_) are consistently defined.

#### Model evaluation

The neural network model performs larval pose prediction orders of magnitude faster than the template-based pose estimation. Pose prediction using the neural network model takes ~300 milliseconds per frame, significantly faster than the 10-12 minutes per frame for template-based pose estimation (**Materials and Methods(b) Template-based pose estimation**). Thus, pose prediction using a neural network model is a much more practical approach compared to the template-based approach. We next asked how the pose prediction accuracy of the neural network’s model compares to that of the template-based approach, using a quantitative assessment of the accuracy of the two approaches.

To evaluate our neural network model, we rendered the physical model images using the inferred pose and compare them to the corresponding three real input images. We computed the Pearson’s correlation coefficient between each of the three pairs of input and physical model images. The network’s pose prediction score for a given frame is defined as the minimum of the three correlation coefficients. Before computing the correlation coefficients, a binary mask was obtained by segmenting the physical model images. The correlation coefficient was only calculated for non-zero pixel locations occupied by this mask. Since the input images have a noisy background and the physical model images have a black background, there is no correlation between the background pixels of the pair of images. Using a mask ensures that the value of the correlation coefficient is determined mainly by the pixels comprising the images of the larval body, and not those that make up the background. **Figure 2a** shows the network’s pose prediction score on our entire free swimming, acoustic startle, and dark flash dataset comprising 630 movies (68181 frames). The quality of the pose estimate can be interpreted by comparing the physical model image and the recorded image for different values of the correlation coefficient (see **Figure 2b**).

**Figure 2:**
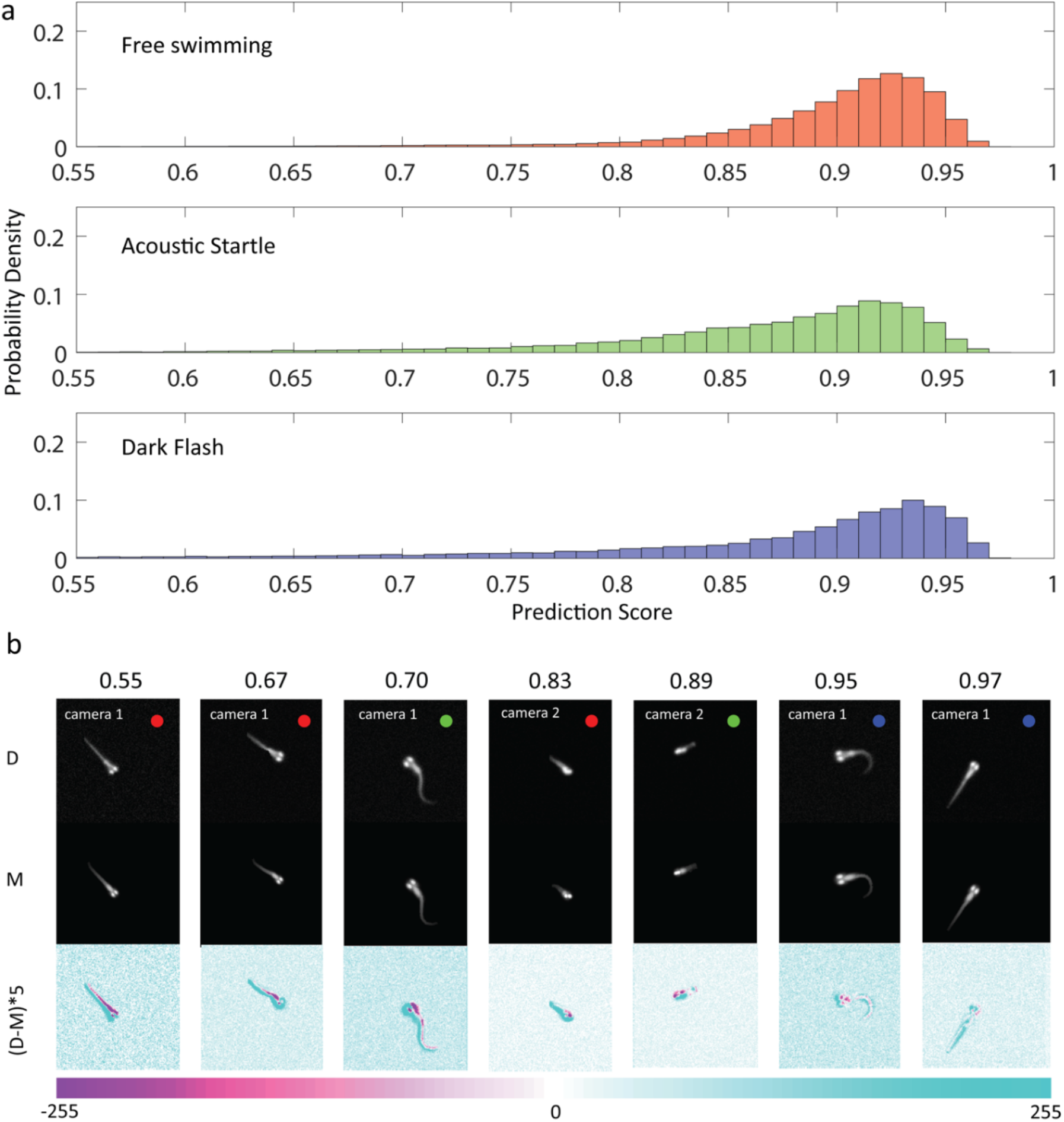
Evaluation of the predicted poses on real data. (a) Histograms of the pose prediction scores shown for all three experimental contexts (red, green, blue) comprising 630 movies (68181 frames). The pose prediction score is defined as the minimum of the correlation coefficients between the recorded image and the physical model image computed independently for the three camera views. Pose prediction was performed on images collected from the three experiments separately. (b) Comparison between experimental and model images representative of the prediction scores in (a). Only one of the three views of the fish is shown. D: experimental Data collected with the three-camera system, M: Model images rendered using the voxel-based model. (D-M)*5: Difference image of the experimental Data and Model, amplified by a factor of 5. Pixel values in the range [-255,255] of the amplified difference image can be interpreted by the colorbar below. Colored dots (red, green, blue) denote the experimental context for each image.

We trained the network with several datasets to evaluate its robustness to variability of the input data. Specifically, we generated three independent training datasets, using (1) 100% (*N* = 35714) and (2) 25% (*N* = 8928) of the poses in the ensemble of real poses, and (3) 100% of free swimming poses (*N* = 9536) in the ensemble of real poses. We also evaluated the accuracy of the template-based pose estimations as a comparison. Evaluations were performed over all 424 swim bouts (41756 frames) for which the computationally expensive template-based pose estimation was performed. We find that the trained CNN model performs marginally better (mean pose prediction score = 0.91) than the template-based pose estimation technique (mean pose prediction score = 0.90) on average (**SI Figure 6**). Thus, the neural network model is superior to the template-based pose estimation in speed without sacrificing accuracy. We also find that reducing the amount of experimental data (25% of all poses) used in the training dataset does not deteriorate the performance of the network model (pose prediction score = 0.91). However, using only free swimming poses for generating the training dataset does decrease the network’s performance (mean pose prediction score = 0.90). More specifically, in this case, the pose prediction accuracy is marginally improved for free swimming data and marginally deteriorated in the acoustic startle and dark flash data (**SI Figure 6**).

### 3-D kinematic parameters reveal unique features of larval swims across the three experiments

We use the network predictions to inspect swimming parameters in the 3-D trajectories of larval swims. We performed pose prediction using the neural network over all the video recordings, comprising 630 swim bouts. We rejected any swim bout where at least one frame has a pose prediction score smaller than 0.85, resulting in 478 swim bouts (free swimming – 258, acoustic startle 106, acoustic startle – 114). **Figure 3a,b** shows the center of mass trajectories (*x*_0_, *y*_0_, *z*_0_) of the larvae recorded in free swimming, acoustic startle, and dark flash experiments. We find that these trajectories are symmetric in the *x_lab_-y_lab_* plane. Thus, we rotated each trace along the *z_lab_*-axis such that the initial headings in the *x_lba_-y_lab_* plane, *θ*_0_ (*t* = 0), are oriented along the -*x_lab_* direction. As seen in **Figure 3b**, the vertical motion of the larva has a significant component, with z-displacements several times larger in magnitude than the larva’s body length and the 2-3 mm height typically provided in constrained 2-D experiments (6,21,23). During free swimming (red), vertical displacements span a range from −2.4 mm to 6.8 mm (*N* = 258), mainly characterized by ascents. In contrast, in acoustic startle (green) and dark flash experiments (blue), vertical displacements range from −3.9 to 3.4 mm and from −4.7 to 0.7 mm, respectively, and are mainly characterized by descents.

**Figure 3:**
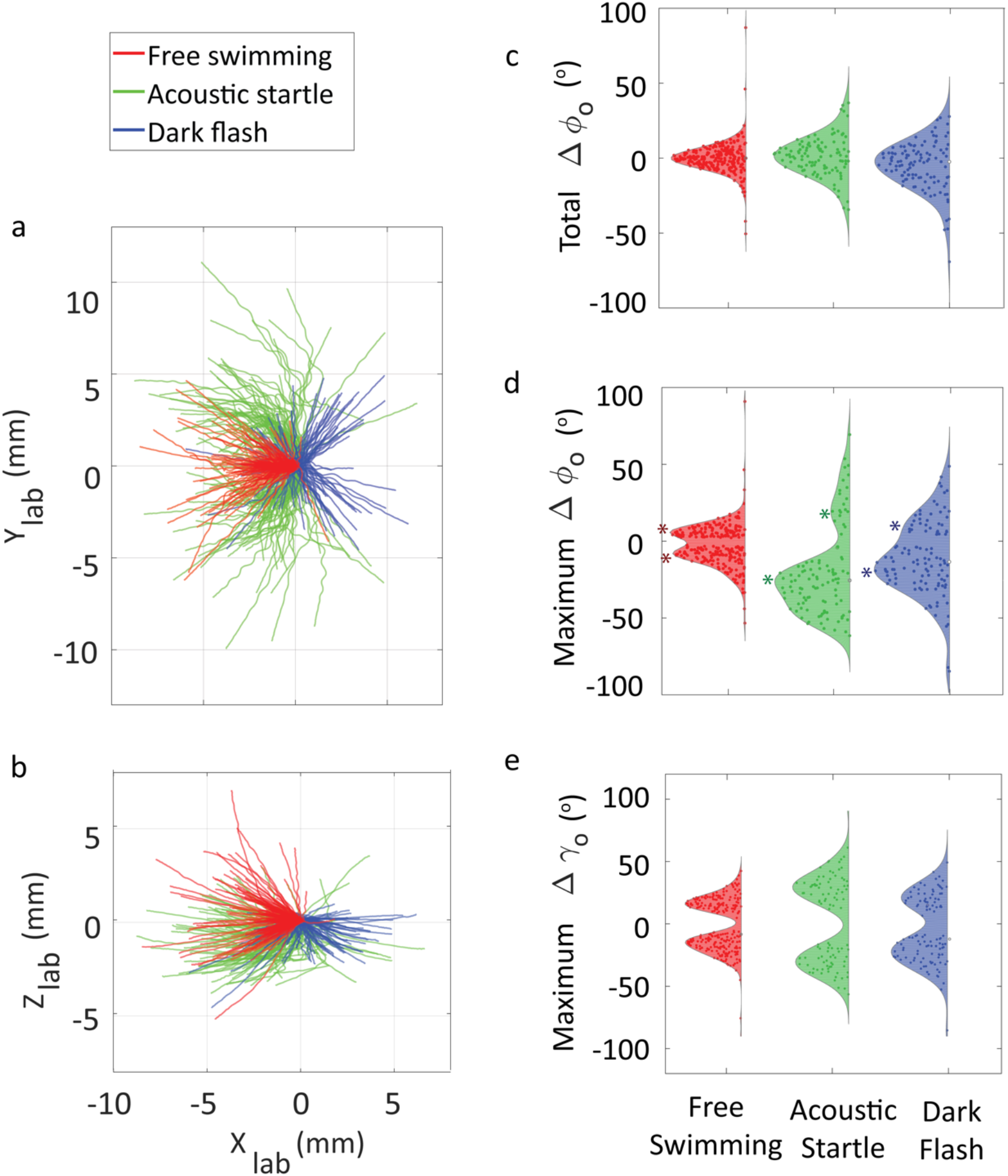
3-D kinematic parameters reveal unique features of larval swims across the three experiments. (a) Trajectories of the center of mass of larvae swimming in a 3-D environment (*N* = 478) in the *x*_lab_-*y*_lab_ plane, in all three experimental contexts (red, green, and blue). (b) 3-D trajectories show a significant z-component, larger than the depth of 2-3 mm in constrained 2-D setups. (c) Distributions of the total change in inclination angle *φ*_0_ over a swim bout. The distributions in all three experiments peak at Δ*φ*_0_ = 0, showing that the larvae tend to reorient themselves along the same vertical direction at bout termination as before the initiation of the swim bout. (d-e) Distributions of the maximum change in inclination angle Δ*φ*_0_ (d) and roll angle Δ*γ*_0_ (e) over a swim bout. Escape responses evoked by acoustic startles (green) involve the largest component of these motions made accessible exclusively in a 3-D measurement. The asymmetry in the distributions of max(Δ*φ*_0_) of acoustic startle and dark flash responses in (d) is visualized by differences in the peaks of their distributions above and below max(Δ*φ*_0_) = 0 (marked by *). This asymmetry shows the propensity of the larvae to dive in these experiments.

We also analyzed the inclination (*φ*_0_) and roll angles (*γ*_0_) of the larvae, parameters inaccessible in 2-D experiments. We calculate the range of angles in these distributions by fitting a kernel density estimate and finding the angles between which the cumulative distribution function lies within 0.01 and 0.99. We find that *φ*_0_ varies between −34° and 49° in free swimming experiments, −42° and 40° in acoustic startle experiments and −41° and 42° in dark flash experiments. *γ*_0_ varies between −24° and 17° for free swimming experiments, −26° and 23° for acoustic startle experiments, and −24° and 22° in dark flash experiments. The distribution of total change in inclination *φ*_0_ over a swim bout is shown in **Figure 3c**. We find that in a majority of swim bouts, larval inclination angles return to their initial value. **Figure 3d-e** show the distribution of maximum change in inclination max(Δ*φ*_0_) and roll max(Δ*γ*_0_) in a swim bout. We find that max(Δ*φ*_0_) varies in the range of −34° to 31 ° in free swimming experiments, −66° to 61 ° in acoustic startle experiments and −79° to 45° in a dark flash experiment. The asymmetry between negative and positive max(Δ*φ*_0_) (**Figure 3d**) for acoustic startle and dark flash trajectories is consistent with the preponderance of dives in those experiments. On the other hand, the corresponding distribution of roll angles max(Δ*γ*_0_) is roughly symmetric, with the larvae exhibiting a larger range of rolls in response to acoustic startle stimuli (range for free swimming: −40° to 37°, acoustic startle: −63° to 74°, dark flash: −55° to 53°; see **Figure 3e**).

## Discussion

The reconstruction of larval zebrafish swimming behavior in three dimensions has long been a challenge, with several obstacles such as the high speed and small size of the animal, refraction from water and glass, and the transparent body of the larvae. While the latter trait facilitates imaging of its organs and tissues (25,36,37), it complicates the imaging of behavior because the posterior part of the tail is more difficult to distinguish from the background. Here, we present a tracking algorithm based on a physical-model trained neural network analysis of fish videos that produces an accurate coordinate representation of its poses and behavior in 3-D using relatively low-resolution cameras (648×488 pixels) taken by an affordable multi-camera setup. Its robustness is illustrated by the quantitative evaluations of the network’s performance. We find that zebrafish larvae swimming in an unrestrained 3-D space perform significant vertical motion and rolls, features either absent or not detected in classical 2-D experiments. We anticipate that with more sophisticated cameras, the system may in the future allow the study of fins, movement of eyes (38), and other attributes, which can be incorporated easily into the physical model by adding a few additional coordinates.

The approach of synthetically generating digital images to tackle several pattern recognition problems using a neural network has been used in the past (12,39–46). It provides two advantages over collecting and manually annotating real training data: (1) generation of an arbitrarily large dataset to train artificial neural network models, and (2) accurate annotations of the dataset without needing human intervention. However, in many of these examples, features of the digitally generated images are significantly different from those of real images. This necessitates developing new techniques for integration of digitally generated data into the analysis pipeline. Research in the field of *domain adaptation* is an ongoing effort to address these challenges (42,47–49). A few notable exceptions where a machine learning model trained on digitally generated images can be directly used on real images involve the explicit use of real images to generate the training dataset (12,50). Here, simple transformations (like scaling, rotation, translation) applied to real images are used to generate new training examples. Such a use of real images plausibly preserves the relevant image features in the generated training dataset, obviating the need for any other modifications downstream to tune the network to perform with sufficient accuracy on real data. However, the use of such image transforms is not practical in our context due to reasons ranging from self-occlusion to the non-trivial coupling of information between the three input images. We thus render the physical model images independently using a physical model-based approach without explicitly using any real images. Our digitally generated data seamlessly integrates into the training pipeline.

We anticipate that this rich and annotated digitally generated dataset can be used in many other applications. One such immediate application may be studying a school of larval zebrafish. Our current network model only predicts the pose of a single larva. Research groups have already had success tracking multiple animals using artificial neural networks (51–53). A digitally generated dataset of a school of larval fish generated using the physical model could in principle be used to train a network inspired from one of these studies, opening avenues to questions about the social behavior of zebrafish larvae freely swimming in a 3-D environment. The physical model images may also be used to generate a training dataset for larval pose estimation in classical 2-D experiments where a single overhead camera is used for imaging. 2-D pose estimation using such a dataset may be performed using an alternate, simpler neural network model or any of the existing neural network architectures that have shown promising results for 2-D animal pose estimation (10,12,16,53). Our fully annotated larval zebrafish dataset can also serve as a benchmark for upcoming tools aimed not only at animal pose estimation, but more generally to computer vision problems like 3-D reconstruction and pose estimation.

Since the parameters of the projection function are not baked into the network model, the convolutional neural network model is agnostic to camera calibration. We in fact find that the network’s performance does not change due to small drifts in the cameras occurring over time. This is reflected in the fact that the projection function used to generate the training dataset was different from that calibrated for free swimming experiments. However, we see that the pose prediction score on free swimming swim bouts is consistently higher for all the network models that we tested (**SI Figure 6**).

Our dataset of temporally independent poses may also be used to generate one of temporally correlated sequences of poses, as exhibited in stereotyped behaviors. This dataset may be useful to learn an algorithm that can predict the future poses of a larvae, given a sequence of poses in the past. Such an algorithm could be useful in improving hardware that requires online tracking of fast swimming larvae (25) for simultaneous behavioral/neuronal recordings. The use of a generative model that learns temporal relationships in the data may help create such a dataset. Such a model may be trained by using a powerful neural network model (54). One could also use a more physics-based approach by learning the postural dynamics using closed-form expressions. An example of such an approach is a recently published work by (4), where high-dimensional postural dynamics of a moving worm were approximated as a series of linear stochastic differential equations, inferred from the data.

We also anticipate that our dataset of 3-D larval poses will rouse interest among ethologists studying zebrafish larvae to investigate novel aspects of their 3-D poses and dynamics, previously unexplored in constrained 2-D experiments. We hope that our system significantly reduces the challenges for zebrafish researchers to perform more behavioral experiments with zebrafish larvae in a native 3-D environment.

## Materials and Methods

### (a) Instrumentation

#### Animals

Adult AB genotype zebrafish larvae were obtained from the Zebrafish International Resource Center (ZIRC, Oregon). Larvae used in the experiments were obtained by breeding the adult fish in the lab and raised at 28°C in Danieau’s solution (31) until 6 dpf (days post fertilization). All behavioral experiments were performed on larvae between 6 and 9 dpf in accordance with the protocol (#19100) approved by the Illinois Institutional Animal Care and Use Committee.

#### Experimental setup and data collection

The experimental setups used for the 3-D experiments are shown in **Figure 1—Figure supplement 2**. Fish swimming measurements in 3-D were carried out in a custom-built cubic glass tank of dimensions 7 x 7 x 7 cm. Zebrafish larvae at 6-9 dpf and measuring a few mm in length were placed in the tank and were allowed to swim freely without external stimulus. Movies of the larvae were obtained at 500 fps using 3 synchronized high-speed cameras (Ximea xiQ MQ003MG-CM) aligned with their principal axes orthogonal to each other and intersecting at a unique point in the center of the tank. The cameras were aligned as follows. Briefly, a mirror was placed facing the camera to be aligned, with its surface along the nearest face of the cubic tank. The center of the camera lens seen in its reflected image was detected using image analysis. In order to align the principal axis perpendicular to the mirror (and hence to the face of the tank), the (detected) center of the lens was made to coincide with the center of the image by rotating the camera on its mount. Similarly, the other two cameras were also rotated. To ensure that their principal axes intersected at the center of the tank, we translated the cameras using the detected centers as reference. Infrared illumination at wavelengths >700 nm enabled the same imaging modality for all experiments, including dark flashes. We used 3 orthogonally placed and diffused infrared lamps (Chanzon, *λ* = 850 nm) with a high-power output (50 W) to compensate for the high frame rate and small optical aperture (large *f*-ratio) of the cameras, which was necessary in order to achieve a large depth of field.

Zebrafish larvae were habituated in the 3-D arena for 10 minutes after being transferred from the incubator. Movies of larval swimming were then collected in three experimental contexts: free swimming, acoustic startle, and dark flash stimulus. Acoustic startles were generated by releasing a heavy cylindrical weight from an electromagnet on the platform supporting the glass cube (**SI Figure 2c**). A dark flash stimulus was provided by initially habituating the larva to a white light LED for at least 4 minutes before instantaneously turning the light off for 10 s (**SI Figure 2b**). Automated stimuli were generated using an Arduino UNO board interfaced with MATLAB and the stimulus source (electromagnet/white LED) triggered using an NPN transistor.

It was rare for a larva to be in the field of view of all three cameras at the same time. Thus, a search for such a larva was conducted once every second and a stimulus was triggered only when one was detected simultaneously by all the three cameras. Since the camera parameters that project a real 3-D point in the 2-D pixel space are non-linear, triangulating the position of the larva using the center of mass detected by the cameras required non-linear optimization. This process is computationally expensive in MATLAB and prevents monitoring the larvae in real time, once a second. Thus, we used the following algorithm to perform this search:

1. Detect the centroids {*C_α_^i^*}, where 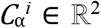 is the centroid of the *i*^th^ larval projection on camera α, where α ∈ {1,2,3} and *i* ∈ {1,2,3,..,*N*} and *N* is the total number of larval projections detected in camera α’s field of view.
2. Determine *R_α_^i^*, the ray of light incident on pixel *C_α_^i^* of the camera sensor array, using Lookup Table B (see **Materials and Methods(a) Instrumentation:** *Camera calibration*: *Reverse mapping from camera coordinates to lab frame via Lookup Table R*). By construction of the lookup table, this ray of light should have originated from a point coinciding with a larva in the tank.
3. Determine all the tuples (*R*_1_^i^, *R*_2_^j^, *R*_3_^*k*^) iterating over all *i, j, k* in the three cameras.
4. For each tuple (*R*_1_^i^, *R*_2_^j^, *R*_3_^*k*^), compute δ = max[*d*(*R*_1_^*i*^, *R*_2_^*j*^), *d*(*R*_2_^*i*^, *R*_3_^*k*^), *d*(*R*_1_^*j*^, *R*_3_^*k*^), where *d*(*R_α_^m^*, *R_β_^n^*) is the minimum distance between the rays projecting on the pixel *C_α_^m^* of camera α and *C*_β_^*n*^ of camera β.
5. If δ < ε, the rays (*R*_1_^i^, *R*_2_^j^, *R*_3_^*k*^) originate from the same larva (ε is a small positive number determined heuristically).

#### Camera calibration: Mapping from lab frame to camera coordinates

In the absence of the water tank, the mapping from a 3-D lab frame coordinate (*x*_lab_, *y*_lab_, *z*_lab_) to a 2-D point on the camera frame (*x_i_*, *y_i_*) is a straightforward linear operation given the camera matrix **P**_i_: [*x_i_*, *y_i_*, 1] = [*x_lab_, y_lab_, z_lab_*, 1]**P**_*i*_ where *i* = {1,2,3} is the index of the camera (55). The camera matrix **P**_*i*_ is a 4 x 3 matrix modeled as a product **M**_*i*_ **K**_*i*_ of a 4 x 3 ‘extrinsic’ matrix **M***_i_* and a 3 x 3 ‘intrinsic matrix’ **K**_*i*_. **M**_*i*_ is determined by extrinsic parameters characterizing the rotation (quantified by the Euler rotation matrix, *R*_3×3_) and translation (characterized by the vector *T*_3×1_) of each camera relative to the lab coordinate system. The ‘intrinsic’ matrix **K**_*i*_ is determined by intrinsic parameters of the camera such as the focal length (in pixels), *f_x_*, *f_y_*; principal point coordinates (in pixels), *μ*_0_ and *v*_0_; and the skew coefficient of the pixels *γ*. A dot grid target (R2L2S3P4, Thorlabs) was used in the camera calibration process to determine all the camera parameters listed in the matrices **M**_*i*_ and **K**_*i*_ (see **SI Table 1**). Several sets of images of the target at different orientations to the camera were taken (shown in **SI Figure 2a**) for camera calibration. The positions of the dots were detected using a custom-written code. Camera calibration was performed with the Computer Vision System Toolbox in MATLAB.

In fish swimming measurements, the projection from a 3-D lab coordinate of a point on the fish to its 2-D coordinate on the camera is no longer a linear operation because of the refraction caused by water and glass. There still exists a function *f_i_* for camera *i* such that (*x_i_*, *y_i_*) = *f_i_*(*x_lab_*, *y_lab_*, *z_lab_*). However, an accurate analytical expression for this function is cumbersome and slow to compute. Rather than seeking the exact solution, we obtained an empirical function *f_i_* by fitting images of the dot grid target. First, we collected a set of pictures of the dot grid target without the cubic tank (**SI Figure 3a**). Then, we placed the filled tank around the target while keeping the target at the same position as before and collected another set of pictures (**SI Figure 3b**). We obtained two sets of 2-D coordinates of the dot patterns, {*r^water^*} and {*r^air^*}, corresponding to the images taken with and without refraction, respectively, where *r^water^* = (*x_i_, y_i_*), and *r^air^* is a vector comprising the *x* and *y* coordinates of the 2-D projection of a dot without refraction (**SI Figure 3c**). The 3-D lab coordinates of the *N* dots {*R_n_^air^*} were then reconstructed from {*r_n_^air^*}, where *n* ∈ {1,2,3,..,*N*} by minimizing the function 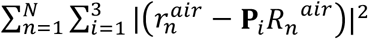. Because the lab coordinate system *x_lab_, y_lab_, z_lab_* in **SI Figure 1c** is based on one of the cameras, with three cameras perpendicular to each other, we found that it was possible to simplify the projection function by calculating the two camera coordinates (*x_i_*, *y_i_*) independently. For instance, for camera 1, in the relation (*x*_1_, *y*_1_) = *f*_1_(*x_lab_, y_lab_, z_iab_*), with *x*_1_ being perpendicular to *y_lab_*, the value of *x*_1_ was independent of *y_lab_*. Thus, the projection function could be separated into two independent functions *x*_1_ = *f*_1*x*_(*y_lab_, z_lab_*) and *y*_1_ = *f*_1*y*_(*x_lab_, z_lab_*), each of which was fit to a cubic function for interpolation to achieve satisfactory accuracy. As shown in **SI Figure 3d**, the fit function *f*_1*x*_ matched the experimental data (plotted as gray dots) well. With the empirical projection functions, we are able to reduce the mean triangulation error from 1.07 mm to 0.03 mm (**SI Figure 3e**), a small fraction of the body diameter of the animal.

#### Camera calibration: Reverse mapping from camera coordinates to lab frame via Lookup Table R

Having found the mapping from 3-D lab frame coordinate (*x*_lab_, *y*_lab_, *z*_lab_) to 2-D point on the camera frame (*x_i_*, *y_i_*), it is necessary for analyzing larval swimming movies to construct the reverse mapping from (*x_i_*, *y_i_*) to the set of 3-D coordinates that project onto (*x_i_*, *y_i_*). Given that the laws of ray optics hold true for our setup, we expect a set of coordinates on a unique ray, **a** + Λ**r**, to be projected, after refraction through the water-glass-air interface, onto a given pixel (*x_i_*, *y_i_*) of the sensor array of camera *i* (**SI Figure 4**). Here, Λ parametrizes the ray, **a** ∈ *R*^3^ defines the origin of the ray in the lab reference frame at Λ = 0 and **r** ∈ *R*^3^ is the directional vector of the line, defining the slopes along the 3 lab coordinate axes. Λ is defined such that Λ = 0 corresponds to the point of intersection of the ray with the plane perpendicular to the principal axis of the camera, passing through the center of the tank. Since determining the set of 3-D points whose projection falls on the pixel (*x_i_*, *y_i_*) involves nonlinear optimization, it is computationally inefficient and thus motivates the use of a lookup table. For each camera, an independent ‘Lookup Table R’ for aiding in reverse mapping stores the vectors **a** and **r** for every pixel (*x_i_*, *y_i_*) of the camera sensor array. The values of **a** and **r** are determined by finding three random 3-D points in the tank that are projected onto the given pixel (*x_i_*, *y_i_*), using the non-linear mapping functions *f_i_*(*x_lab_, y_lab_, z_lab_*) above, and fitting a line through these three points. Lookup Table R is used in triangulating the center of mass of larvae in the tank (see pseudo code in **Materials and Methods(a) Instrumentation:** *Experimental Setup and Data Collection* above) and in fitting the three camera images to a physical model of the larva (see **Materials and Methods(b) Template-based pose estimation:** *Physical Model of the Larva*).

### (b) Template-based pose estimation

#### Preprocessing

The videos of larval swimming were preprocessed to improve the speed and accuracy of larval pose estimates. The raw images were background subtracted to increase the contrast between the foreground and the background and to reduce noise. The background image was created by computing the 90^th^ percentile intensity of every pixel over the entire video. This image was then passed through a Gaussian filter (window size: 5 x 5 pixels, standard deviation: σ = 1) to reduce high frequency noise further. The images of the larva were then segmented into binary images using a heuristically defined intensity threshold, and a bounding box was defined around the larva. The size of the bounding box was determined so that the largest 2-D projection of the larva (seen when the larva is closest to the camera) could be contained within the box. Since multiple projections of larvae were often detected in several frames, the larva of interest needed to be tracked. Tracking over multiple frames was performed by triangulating the centroid of the larva in 3-D coordinates in the current frame and detecting the nearest neighbor centroid in the following frame. For all the analysis downstream, we used this background subtracted and cropped image. Note that all images illustrated in the figures are these preprocessed images.

#### Physical model of the larva

We defined a physical model of the larva using 22 adjustable parameters (**Figure 1c**) and 20 fixed parameters (see below). We showed previously that 10 segments are sufficient to characterize the variations in the shape of the larval backbone (23). Using 10 segments also allows for direct comparison of our results with the previous 2-D data (23). The model is based on a scaffold comprising a chain of 9 rigid segments, allowing for 2 degrees of freedom for every joint connecting successive pairs of segments. Thus, the larval pose, representing the shape of its backbone, is defined in our framework by 16 parameters Δ*θ*_1_,… Δ*θ*_8_, Δ*φ*_1_,… Δ*φ*_8_ that represent the radial and azimuthal angles of each segment measured relative to the larval head orientation. Our choice of the number of segments defining the physical model is constrained only by computational power. In principle, the entire analysis can be reproduced for an arbitrarily large number of segments. The remaining six degrees of freedom—coordinates of the center of mass (*x*_0_, *y*_0_, *z*_0_) and the yaw *θ*_0_, pitch *φ*_0_, and roll *γ*_0_ of the larval head—establish the position and orientation of the larva in the lab reference frame, which is defined with respect to the bottom camera.

The anterior of the larva (eyes, head, and belly) is modeled as a set of ellipsoids, with the first rigid segment used as a scaffold (**SI Figure 5a**). We use the following 19 fixed parameters to define these parts of the larval anterior: brightness of the eyes, head, and belly (3 parameters); distance between the eyes (1 parameter); width, length, and height of the eyes, head, and belly (9 parameters); position of the eyes, head, and belly with respect to the scaffold segment (6 parameters). The position of the belly and head with respect to the scaffold segment can be defined uniquely using only two parameters each, since the belly and head have a constraint to be laterally symmetric about the scaffold segment. The position of the eyes is also defined using 2 parameters, because the centroids of the eyes are constrained on a segment perpendicular to the scaffold segment, with each centroid located at the end of the segment. The length of this segment is twice the distance between the eyes. For a given roll angle of the larva, this segment can be uniquely determined using two parameters: its perpendicular distance from the scaffold segment and its position along the scaffold’s length. The length of the larva *L* brings the count of fixed parameter to 20. *L* also serves to scale the absolute size of the eyes, head, and belly of the physical model of the larva during the second round of optimization (see **Materials and Methods(b) Template-based pose estimation:** *Optimization*). A scaling parameter for the length of the larval anterior, *l_A_*, which depends on *L* and the position of the larva in the tank, is used for rendering 2-D projections of the anterior in the first round of optimization (see **Materials and Methods(b) Template-based pose estimation:** *Optimization*).

The larval posterior (tail) segments are constructed on the scaffold of the remaining 8 rigid segments, with a single fixed parameter scaling the width of the tail segments. This parameter was determined by averaging images of multiple larvae. Since our results were found to be robust to small changes to the width of the tail segments, the scaling parameter for the width is determined uniquely for all swim bouts and kept fixed for the entire analysis. In addition, the length of each tail segment is set to 1/9^th^ times the length of the whole larva *L*, allowing it to vary between larvae.

The larval anterior (head, eyes, and belly) and the posterior (tail segments) are rendered independently. For the anterior, we used a voxel-based 3-D model to generate 2-D projections that recreate the preprocessed images (**SI Figure 1f** and **SI Figure 5a**). The intensity profile *V*(*x_lab_*, *y_lab_*, *Z_lab_*) of the voxels is given by max(**N**(*μ*_eye1_,∑_eye1_), **N**(*μ*_eye2_,∑_eye2_), **N**(*μ*_head_,∑_head_), **N**(*μ*_belly_,∑_belly_)), where **N**(*μ*,∑) is the normal distribution with mean μ and covariance matrix Σ, uniquely defined by the fixed parameters. *μ_foo_* ∈ *R*^3^ is the center of mass of organ ‘*foo*’ and ∑_*foo*_ is a 3 x 3 covariance matrix that encodes the dimensions of *foo*. To obtain 2-D projections of the voxels, we either perform an orthographic projection or a nonlinear projection using the camera matrix, depending on the round of optimization (see ***Image Analysis**: Optimization* and **Figure 1 – Figure Supplement 1d-f**). Pixel intensities of the projections are obtained by summing over the intensity of all the voxels that are projected on the respective pixel. For example, for the orthographic projection with respect to the bottom camera (camera 1 in **SI Figure 1**), pixel intensities *P*_zlab_(*x*_lab_,*y*_lab_) are given by ∑*_zlab_ V*(*x_lab_, y_lab_, z_lab_*).

For the larval posterior, the 2-D projections of the tail segments are rendered using a voxel-equivalent model to make the optimization more efficient in time and memory. In this model, the projections of the tail are constructed on the scaffold of the 8 rigid segments using a series of trapeziums connected by disks (**SI Figure 5b**). The intensity of each of these segments has a Gaussian profile along the width and a linear profile along the length, decreasing in the belly-tail direction. This construction is equivalent to the projection of a voxel-based model consisting of tapered cylinders connected by spheres.

#### Rendering larval projections

We generate realistic larval projections using either of two approaches differing in computational complexity: (i) Using Lookup Table P to render larval projections completely, (ii) Using Lookup Table P to render the larval posterior (tail segments), while the larval anterior is rendered by projecting individual voxels of the voxel-based model (see **Materials and Methods(b) Template-based pose estimation**: *Physical model of the larva*) using the projection function (see **Materials and Methods(a) Instrumentation**: *Camera calibration: Mapping from lab to camera coordinates*). We employ (i) during the first round of optimization (see **Materials and Methods(b) Template-based pose estimation**: *Optimization*), where it is critical to render larval projections rapidly. On the other hand, (ii) is used during the second round of optimization and while rendering physical model images used to train the neural network (see **Figure 1e**), where generating accurate projections is critical while sacrificing on the computational speed.

In order to construct larval projections from Lookup Table P for a given parameter vector **p** ∈ *R*^22^, along with *L*, we render the larval anterior and posterior independently. The procedure for generating digitized larval projections during optimization using Lookup Table P involves determining the appropriate lookup table indices corresponding to the scaling parameter and orientation of the larval anterior and tail segments. The procedure to render larval anterior using Lookup Table P is described in detail in SI: **Rendering larval anterior using Lookup Table P**).

For rendering the tail, this procedure is not necessary because the indices of the tail segments, by construction of Lookup Table P for tail segments, are defined with respect to the orientation, length, and offset of the 2-D projections explicitly. We first compute the 2-D projections of the larval backbone (the chain of 8 rigid segments forming the posterior) using the non-linear projection functions *f_i_*(*x_lab_, y_lab_, z_lab_*) described above (see **Materials and Methods(a) Instrumentation**: *Camera calibration: Mapping from lab to camera coordinates*) and determine their projected length. This gives the index corresponding to the scaling parameter *l_T_*. The entry of Lookup Table P determining the orientation for the tail segments can be evaluated directly by computing the orientation of the 2-D projection of the individual tail segments forming the larval backbone. Lastly, the subpixel offsets can be determined by computing the fractional part of their *x* and *y* coordinates.

We also render the larval anterior directly using the voxel-based model in some cases, as described above in this section in (ii). Rendering every voxel individually using the non-linear projection function (see **Materials and Methods(a) Instrumentation**: *Camera calibration: Mapping from lab to camera coordinates*) assures higher accuracy but is computationally expensive. While rendering the larval anterior in this way, a voxel-based model of the 3-D larval anterior is first generated (see **Materials and Methods(b) Template based pose estimation**: *Physical model of the larva*). Every voxel is mapped from the 3-D lab-coordinates to the 2-D camera coordinates and the voxel intensities projected at every camera coordinate are summed, resulting in a 2-D image. Further, the intensity of the 2-D image is scaled such that the brightest pixel has a value of 255.

#### Optimization

The physical model of each larva is constructed from the unique chain defined by the parameter vector **p** ∈ R^22^: **p** = (*x*_0_, *y*_0_, *z*_0_, *θ*_0_, Δ*θ*_1_,… Δ*θ*_8_, *φ*_0_, Δ*φ*_1_,… Δ*φ*_8_, *γ*_0_), in addition to the total length *L* of the larva. In order to compute the larval pose in any frame of a swim bout movie, we perform a search over the space of 22 adjustable parameters such that digitally generated larval projection images of the physical model best match the three views recorded by the cameras. A cost function, defined as the sum of the 2-norms of the three difference images (of the background subtracted image and the digitally generated projection) for the three cameras, is minimized globally using the *patternsearch* algorithm in MATLAB’s Global Optimization Toolbox. Each frame of the bout is analyzed in parallel on a node of an Intel Xeon Gold 6150 @2.7 GHz cluster.

The entire pipeline for the analysis of swimming movies is described in **SI Figure 1**. We performed the optimization of the physical model against the collected data in two rounds. In the first round, a coarse estimate of the model’s parameter vector **p** is generated using a lookup table (see **Materials and Methods(b) Template-based pose estimation:** *Lookup Table P*) that stores a large array of larval projections (**SI Figure 1e**). In the second round, fine optimization improves the estimate in a small neighborhood of **p** obtained in the first round (**SI Figure 1f**). Here, the larval projections are obtained by rendering the voxels defining the physical model at every iteration (see **Materials and Methods(b) Template-based pose estimation:** *Rendering larval projections*).

Before initializing the optimization, we estimate the fixed parameter *L* from the length of the binary, segmented image of the larva from the bottom camera in the initial and final 5 frames of the swim bout, during which the larval backbone is expected to be straight. Specifically, we first find the best orientation angles of the larva (*θ*_0_, *φ*_0_) that minimize the cost function, assuming a physical model with a straight backbone and a larval length of 4.5 mm. *L* is averaged over all 10 frames.

In the first iteration of the coarse optimization, the adjustable parameters are initialized by an approximate guess of the centroid of the larva (*x*_0_, *y*_0_, *z*_0_), triangulated using the centroids of the segmented larval images captured in the three cameras. This triangulation is performed using the nonlinear projection function by nonlinear optimization (*fmincon* function in MATLAB). Since recording is started before the stimulus is generated, we expect the larval backbone to be a straight line in the first frame and find the best orientation (*θ*_0_, *φ*_0_, *γ*_0_) that minimizes the cost function. For this step, we use a brute force approach by iterating over a large number of orientations (*θ*_0_, *φ*_0_, *γ*_0_) and render the corresponding digital projection of the larva (see **Materials and Methods(b) Template-based pose estimation:** *Rendering larval projections*). The constraint on the shape of the backbone significantly reduces the volume of the search space and makes the brute force approach tractable. For the first frame, where the larval backbone is straight, we use the parameter vector obtained from the brute force approach as an initial guess and compute the parameter vector **p**_0_ that minimizes the cost function, using the *patternsearch* algorithm in MATLAB. For all the succeeding frames, the initial guess is the optimized parameter vector from the preceding frame.

In the succeeding iterations of coarse optimization, Lookup Table P is used to render the larval anterior and the larval posterior, given the parameter vector **p** (see **Materials and Methods(b) Template-based pose estimation***: Rendering larval projections*). During rendering, for pixels shared by the projection of overlapping larval anterior and posterior segments, the higher pixel intensity value is selected. The estimate **p**_0_ is computed by optimization of the cost function described previously. Since the Lookup Table P entries are rendered for discrete orientations of the larva, the estimate of **p**_0_ is coarse grained, and we find that the estimates of roll angle *γ*_0_ are often inaccurate. We thus pass **p**_0_ through a round of fine optimization in the local neighborhood of **p**_0_.

During fine optimization, we use **p**_0_ as the initial guess and search for local minima in a small neighborhood around this guess. Here, an image of the larval anterior is rendered by performing a refraction-based non-linear projection of the voxel-based model for every iteration (see **Materials and Methods(b) Template-based pose estimation:** *Rendering larval projections*). Moreover, the optimization is performed for the anterior and the posterior independently to reduce the volume of the search space. In order to check for convergence, we then perform a final iteration of optimization over the 22-dimensional space and optimize the anterior and the posterior together. Since fine optimization is extremely slow (10-12 minutes /CPU/frame), we manually selected ~150 swim bouts each from free swimming, escape response and dark flash experiments for this task. The fits obtained from coarse optimization were visually inspected to select the 450 swim bouts.

### (c) Neural network pose estimation

#### Generation of ensemble of physical model poses

A total of 35714 real larval poses were selected to generate the ensemble of physical model poses (**Figure 1c**), later used to generate the training dataset. These real larval poses are a subset of the poses estimated using the template-based approach (**Materials and Methods(b) Template-based pose estimation**) that have a pose prediction accuracy greater than or equal to 0.9. We found that the set of larval poses was highly skewed, favoring poses with a nearly straight backbone (**Figure 1b** *Biased data*). Thus, before we could use these larval poses to generate the training dataset, we sought to resample the larval poses uniformly.

First, a kernel density estimate was performed using a Gaussian kernel to approximately infer the probability distribution of larval poses in the (<|Δ*θ*_i_|>, <|Δ*φ*_i_|>) space. In order to evaluate a reasonable bandwidth for the Gaussian kernel, we performed a search over 50 bandwidths uniformly selected in the range [0.01 rad, 0.1 rad]. The data were iteratively split into a test set and a training set in a 1:19 ratio. A kernel density estimate was evaluated using the training set and the iterable bandwidth. For each iteration, the log likelihood of the test set was calculated. The bandwidth that maximized the log likelihood of the test set was selected. The search for bandwidth was implemented using the *GridSearchCV* function of python’s scikit-learn (version 0.24.2) library. A 20-fold cross-validation was used to iteratively split the ensemble of real poses (**Figure 1a**) into test set and training set. The probability distribution 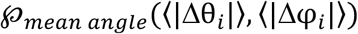 computed using the kernel density estimate was used to sample 2500 poses that are uniformly distributed in the (<|Δ*θ*_i_|>, <|Δ*φ*_i_|>) space (**Figure 1b** *Uniform data*). To achieve this uniform sampling, we choose 2500 poses, each with a probability inversely proportional to 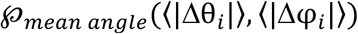.

Next, we generated an ensemble of 500,000 physical model poses, which are modelled on real larval poses. Using the 2500 poses in the uniform data (**Figure 1b** *Uniform data*), we computed their probability distribution 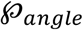 in the 18-dimensional space defined by Δ*θ*_1_… Δ*θ*_8_, Δ*φ*_1_… Δ*φ*_8_, *φ*_0_, *γ*_0_. The kernel bandwidth used to compute 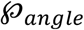 was inferred using the same procedure as described in the previous paragraph. We sampled 500,000 vectors conditional on 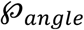 and to each vector, assigned *x*_0_, *y*_0_, *z*_0_ and *θ*_0_ sampled from a uniform distribution. The range of (*x*_0_, *y*_0_, *z*_0_) was constrained by the dimensions of the imaging volume in the fish tank, while the range of *θ*_0_ was chosen as (−*π, π*].

#### Preprocessing training dataset

Using the ensemble of physical model poses, we generated a training dataset of 500,000 physical model images to train the neural network. Generation of all the physical model images was performed in 7 hours using MATLAB on 25 Xenon Gold 6150 CPUs. For every physical model pose, a set of three physical model images was rendered, as seen by the three cameras in our setup (**Figure 1d,e**), using the voxel based model (see **SI Figure 1f** and **Materials and Methods(b) Template-based pose estimation**: *Physical model*) and the projection function (**SI Figure 3**). All the physical model images were then pre-processed before passing to the neural network.

More specifically, the three 2-D larval projections corresponding to each of the 500,000 physical model poses were rendered onto three 141×141 images with pixel values scaled between 0 and 1. The larval projections were displaced at random by 0-20 pixels, both horizontally and vertically. To each image, we added Gaussian noise, whose mean and standard deviation were sampled randomly in order to mimic the background noise in the real images. The brightness at each background pixel of the projections of the bottom camera and side cameras is sampled from Gaussian distributions *N*(*μ_bottom_, σ_bottom_*) and *N*(*μ_side_, σ_side_*), respectively. For each projection of the bottom camera, the mean *μ_bottom_* of this Gaussian distribution is a product of two random variables— *X_μ,bottom_*, drawn from a uniform distribution between 0 and 1/255 and *Y_μ,bottom_*, drawn from a Gaussian distribution with mean 50 and standard deviation 10, while the variance, *σ*^2^_*bottom*_ of the Gaussian distribution is a uniform random variable between 20/(255)^2^ and 70/(255)^2^. For each projection of the side camera, the mean *μ_side_* of the Gaussian distribution is a product of two random variables—*X_μ,side_*, drawn from a uniform distribution between 0 and 1/255 and *Y_μ,side_*, drawn from a Gaussian distribution with mean 20 and standard deviation 10, while the variance, *σ*^2^_*side*_ of the Gaussian distribution is a uniform random variable between 10/(255)^2^ and 60/(255)^2^. Next, we rescaled the pixel values between 0 and 255, to mimic the format of the recorded images – unsigned 8-bit.

#### Preprocessing real data

The real images were preprocessed before passing as inputs to the neural network. We performed background subtraction and cropping on every frame (see **Materials and Methods(b) Template-based pose estimation:** *Preprocessing*) obtained from the recorded videos to validate the trained neural network. The neural network accepts three images each of dimensions 141×141. Since the cropped 2-D larval images were of varying size, they were also padded appropriately with zeros to dimensions 141×141. In order to simulate background noise in the padded region, we first generated a background vector of pixel values along the pre-processed image’s border. This border was defined by the region with 5 pixels of the image boundary. The padded region was assigned pixel values from this vector, selected uniformly at random. This procedure of simulating the background noise works in most cases because the background distribution is uniform, i.e. the distribution of the intensity in the border is the same as that in any other region of the image not occupied by the larva. In some frames, there is a possibility of an irrelevant larva (from the context of pose estimation) occurring in the 5-pixel border. In order to avoid including pixels corresponding to a larva in the background vector, we excluded all pixels that have a value greater than 3σ with respect to the background vector’s mean. Notably, this procedure is not ensured a 100% success in simulating the background. We find rare cases where the padded region’s noise distribution does not represent the background noise distribution of the pre-processed image. The network predictions are inaccurate for such frames. One could overcome this issue by originally cropping 141×141 windows around the larva before performing pre-processing. However, such cropping dimensions may not be practical to achieve in cases where the larva occupies a region near the frame’s border. For our network evaluation, we challenge the network model by performing the above padding procedure for all frames, essentially assuming the worst case scenario of the larva always being in the border region.

The comparison of template-based pose estimations and neural network pose estimations (see **SI Figure 6**) was performed over 424 swim bouts (41756 frames), for which the computationally expensive template-based pose estimates were evaluated. For a fair comparison between the template-based pose estimations and the neural network pose estimations (see **SI Figure 6**), we manually discarded all frames that had more than one larva in the field of view (FOV). Since the network model is only trained with a training dataset involving single larvae, we observed that the network predictions are inaccurate when there are multiple larvae in the FOV.

While analyzing our entire dataset of 630 movies (68181 frames) to assess the network’s performance (see **Figure 2**) and larval swimming kinematics (see **Figure 3**), the issue of having multiple larvae in the FOV was resolved by first using a binary mask of the larva of interest and assigning all the background pixels an intensity of 0. Realistic background noise was then artificially rendered by the procedure described above. The larval mask was obtained by using the first round of optimization of the template-based pose estimation (see **Materials and Methods(b) Template-based pose estimation**: *Optimization*), and rendering a physical model image based on the estimated pose. Pose estimation using the first round of optimization takes 2 minutes per frame with reasonably accurate pose estimates. This means that some of the inaccurate larval masks used to remove an irrelevant larvae also remove parts of the larva of interest. Neural network pose predictions corresponding to such cases are inaccurate and are automatically filtered out by setting a threshold on the accepted prediction score for analysis. We choose a threshold of 0.85 for our analysis in **Figure 3**.

Here we emphasize that we do not expect the user to manually discard frames having multiple larvae in the FOV to generate pose estimations. We also do not propose generation of larval masks to remove irrelevant larvae from the FOV. Both these procedures are performed in this cases for a fair assessment of the network’s performance. In order to estimate the larval pose directly from video recordings and only using the neural network, a user may use one or both of the two approaches below:

1. Perform experiments with low larval density, such that frames with multiple larvae in the FOV are rare
2. Automatically discard network outputs for frames with multiple larvae in the FOV based on a cut-off on the pose prediction score

#### Convolutional neural network model

Our network consists of an encoder of four bottleneck residual blocks as implemented in (34) with 32, 64, 128, and 256 channels, respectively. The convolutional operations involved kernel size = 3×3, stride = 1 and padding = 3. The encoder is followed by a decoder, consisting of three fully connected layers of dimensions 1×288, 1×144, and 1×72, respectively. The output of the decoder is passed through a sigmoid layer and scaled between 0 and 141. This scaled output (a 1×72 vector) is reshaped into three 2×12 matrices, corresponding to the 2-D projection of the larval pose on the three camera views. Every convolutional and fully connected layer is preceded by a batch normalization (Ioffe & Szegedy, 2015) layer and succeeded by a leaky rectified linear unit activation function (57) (negative slope = 0.01). The network was trained for 120 epochs using the Adam optimizer with a learning rate of 0.001 and a batch size of 300. The convolutional neural network was implemented using the PyTorch library (version 1.10.1) and trained in parallel on 2 NVIDIA V100 GPUs (58). The network is completely trained in 5-6 hours. Network validation was performed on a single NVIDIA A100 GPU.

The loss function for training the network consists of two terms: backbone loss and eye loss. The backbone loss is simply the root mean squared error between the 10 2-D projection pose coordinate predictions for the three camera views (3×2×10 vector) and the corresponding ground truth. The eye loss is similarly defined by the root mean squared error between the 2 2-D eye coordinate predictions for the three camera views (3×2×2 vector) and the ground truth 2-D eye coordinates, with a small modification: the eye loss is made symmetric with respect to the eye indices. More specifically, the eye loss is defined as:

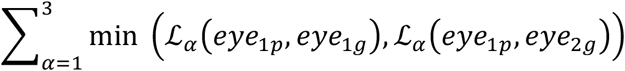

where subscripts 1 and 2 for the eye correspond to the 11^th^ and the 12^th^ coordinate of the 2×12 dimensional 2-D projection pose coordinates, subscripts *p* and *g* for the eye correspond to the network prediction and the ground truth and the 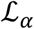 corresponds to the root mean squared error for camera *α*. The total loss function is defined as (backbone loss) + *λ*\*(eye loss), where *λ* captures the relative importance of the eye loss with respect to the backbone loss. For all our neural network experiments, we set *λ* = 5.

#### Triangulation of 3-D pose coordinates

The 10 3-D backbone coordinates are triangulated using the projection function and the 2-D projection pose coordinates obtained as the neural network’s output. Each of the 10 points is triangulated independently by minimizing the least square error between the 2-D coordinate obtained from the neural network and the 2-D projection of the triangulated coordinate obtained using the projection function. We use the center of the tank as the initial guess for this coordinate and implement the optimization using the *least_squares* function of the scipy.optimize package (version 1.7.3) in Python 3.9.9.

The coordinates of the eyes cannot be triangulated directly because of the symmetry in the loss function with respect to the eye coordinates. The symmetry in the loss function causes an ambiguity in the indices of the eyes in the network’s predicted 2-D projection pose coordinates. For example, the network may predict the right-eye as the 11^th^ 2-D coordinate in its 2×12 output vector for the bottom camera, but the 11^th^ coordinate in the output vector for one of the side cameras may be the left-eye. In such a case, one cannot directly triangulate the 11^th^ coordinates in the three views to obtain the 3-D coordinate of one of the eyes. This ambiguity of eye indices is tackled by first permuting over all 8 combinations of eye indices and triangulating the 3-D coordinates of the two eyes in each case. The indices that result in the least triangulation error are accepted. This ambiguity of eye indices is tackled by first permuting over all 8 combinations of eye indices and triangulating the 3-D coordinates of the two eyes in each case. The indices that result in the least triangulation error are accepted.

## Acknowledgements

We thank members of the Chemla and Gruebele laboratories for scientific discussions.

## Supporting information

### Lookup Table P

Lookup Table P of larval projections is used to efficiently render larval projections. Instead of rendering a 3-D digital larva in every iteration of the optimization, we store an array of possible larval projections in a lookup table that can be accessed quickly at every iteration. This lookup table broadly consists of two parts: projections of larval anterior (eyes, head and belly) and projections of larval posterior (tail segments). During the first round of optimization (see **Materials and Methods(b) Template-based pose** estimation: *Optimization*), we use both parts of the lookup table (projections of anterior and posterior) to completely render the larva. During the second round of optimization (see **Materials and Methods(b) Template-based pose** estimation: *Optimization*) and while generating physical model images used to train the neural network model (see **Figure 1e**), only the projections of larval posterior are rendered using Lookup Table P, while the larval anterior is rendered directly from the voxel-based model (see **Materials and Methods(b) Template-based pose estimation**: *Physical model of the larva*).

We note that the 2-D larval projections are a function of both the orientation and position of the larva in the tank. A brute-force construction of the lookup table indexed by the position and orientation of the larva in the tank would require too much storage memory. (For example, if the lookup table stores entries corresponding to the larva located in *ℓ* positions along each direction, the dimensions of the lookup table scales as *ℓ^3^*. Given that a displacement of 0.3 mm in the tank causes a visually recognizable change in the projection and that the field of view is 3 x 3 x 3 cm, *ℓ* has to be at least 10.) Such a lookup table would also generate many degenerate projections. As detailed below, we instead generate a ‘Lookup Table P’ that is indexed by as few variables as possible, and whose size is minimized by exploiting rotational and inversion symmetries.

As with the physical model of the larva, the anterior of the fish (head, gut) and posterior (tail) each have their own Lookup Table P and indexing schemes. For the anterior, each lookup table entry is generated from orthographic projections of the 3-D voxel-based physical model (see **Materials and Methods(b) Template-based pose estimation***: Physical model of the larva;* **SI Figure 1** and **SI Figure 5a**). It is indexed by only 6 variables: the orientation of the larva (*θ*_0_, *φ*_0_, *γ*_0_), a scaling parameter *l*_A_, and sub-pixel offsets (*δ_x_*, *δ_y_*) of the projection. *l*_A_ accounts for the size of the projection, which varies with the position of the larva in the tank. The array of lookup table entries for the bottom and two side cameras are constructed independently to account for the differences in illumination and sizes of larval projections resulting from the bottom camera being closer to the tank compared to the side cameras. Each entry of the lookup table for the bottom camera has a resolution of 49 x 49 pixels and the table has 18 entries of *l*_A_, 180 entries of *θ*_0_ in the range [0, *π*] (entries for the remaining two quadrants can be generated using rotation and/or inversion), 11 entries of *φ*_0_ in the range [0, *π*/2] (entries for the remaining three quadrants can be generated using rotation and/or inversion and correction for subpixel offset), 11 entries of *γ*_0_ in the range [-5*π*/12, 5*π*/12], and 5 entries each for subpixel offsets of 0.2 pixel steps in two orthogonal directions. The lookup table for the side cameras has 18 entries of *l_A_*, 21 entries of *θ*_0_ in the range [0, *π*], 121 entries of *φ*_0_ in the range [-*π*/2, *π*/2] (entries in the remaining quadrants can be generated by rotation and/or inversion), and 5 entries each for subpixel offsets. The step size of the angles is decided such that the length of the 2-D projection of the anterior increases by a constant amount as one iterates over the angles. This ensures that the space of 2-D projections of larval heads is homogeneously sampled. Integral values of pixel offsets are degenerate.

As the 2-D projections of the larval posterior are not rendered using a voxel-based model, Lookup Table P is indexed differently for the posterior than for the anterior. The lookup table stores 2-D projections of tail segments indexed by 5 variables: the segment number, length of the 2-D segment *l_T_* (7 entries), subpixel offsets of 0.2 pixel along the horizontal and vertical direction in the image (5 entries each), and the orientation of the 2-D segment in the plane (360 entries). The tail segment in the physical model does not have a roll angle because the larval tail is approximately cylindrically symmetric. The subtle changes in the tail’s projection that might occur due to torsion are not discernable in our images, constraining the tail segments in our physical model to be torsionally rigid. By construction, the tail segments in Lookup Table P can be accessed directly using the 2-D projection of the larval backbone (see **Materials and Methods(b) Template-based pose estimation:** *Rendering larval projections* below).

### Rendering larval anterior using Lookup Table P

To render the larval anterior from Lookup Table P, we first determine the indices corresponding to the orientation. We note that the angles (*θ*_0_, *φ*_0_, *γ*_0_) corresponding to the indices used during construction of Lookup Table P are not the same as the angles (represented in **p**) defining the orientation of the physical model of the larva, if the 2-D projection of the real larva and the orthographic projection of the voxel-based larva are to match (with the exception of the rare case where the larva is in the center of the tank). This difference occurs because Lookup Table P, in order to increase computational speed (see **SI: Lookup Table P**), is constructed using orthographic projections of the voxel-based model. Thus, as an example, for a larva oriented horizontally facing toward the camera and positioned at a certain height above the principal axis, it should be possible to see the projection of the larval belly and head on the 2-D image projected on the camera. In order for the voxel-based model to produce the same 2-D image rendered by orthographic projection, a small positive pitch angle needs to be introduced to the model. As a result, the indices corresponding to the orientation (*θ*_0_, *φ*_0_, *γ*_0_) need to be adjusted based on the position of the larva in the tank.

To determine the adjusted orientation indices, we assume that all the rays emanating from each voxel of the larval anterior that are incident on their respective pixel in the 2-D projection are parallel. We make this assumption since the size of the larva is negligible compared to its distance from any of the cameras. We further assume that the direction of all these parallel rays is the same as that of the ray CI emanating from the anterior’s center of mass (**SI Figure 5a**). Under these assumptions, the adjusted orientation is determined from rotating the larva such that CI is parallel to the principal axis of the camera of interest. To carry out this rotation, we find an arbitrary point I on ray CI, defined as the intersection of ray CI and the plane perpendicular to the principal axis of the camera and passing through the center of the tank. We then find the elevation angle and the azimuthal angle of CI. We note that a set of five points representing the centroid of the larval eyes, belly, head and the larval centroid uniquely define (*x*_0_, *y*_0_, *z*_0_, *θ*_0_, *φ*_0_, *γ*_0_). These five points are rotated through two Euler rotations such that the azimuthal angle and elevation of CI is zero. This changes the parameters (*x*_0_, *y*_0_, *z*_0_, *θ*_0_, *φ*_0_, *γ*_0_) that the transformed five points represent. An orthographic projection of the voxel-based model that has the same orientation as this new set of (*θ*_0_, *φ*_0_, *γ*_0_) will project and occlude similar parts of the larval anterior as those in the real projection. The appropriate lookup table entry corresponding to the yaw, pitch, and roll (*θ*_0_, *φ*_0_, *γ*_0_) of the reoriented model is then used to represent the projection of the larval anterior. The scaling parameter *l_A_* can be determined using trigonometry, given the 2-D projection (and thus, the length) of the head segment and the orientation of the voxel-based model whose orthographic projection is to be invoked.

**Table 1:** Camera calibration parameters: Intrinsic and extrinsic matrices illustrating the camera parameters used for analysis. The extrinsic matrices **M** shown below, encode the orientation and location of the cameras, where the translation vector T is estimated in mm. The intrinsic matrices **K** encode the intrinsic parameters of the camera in pixels (1 pixel = 7.4 μm). The two matrices were used to analyze fish swimming experiments and had to be calibrated periodically to account for drift in the cameras.

| Camera | Intrinsic Matrix<br>$\mathbf{K} = \begin{bmatrix} f_x & 0 & 0 \\ \gamma & f_y & 0 \\ \mu_0 & \nu_0 & 1 \end{bmatrix}$ | Extrinsic Matrix<br>$\mathbf{M} = \begin{bmatrix} R_{3 \times 3} \\ T_{1 \times 3} \end{bmatrix}$ |
| --- | --- | --- |
| 1 | $\begin{bmatrix} 1143.2 & 0 & 0 \\ 0 & 1139.4 & 0 \\ 327.2 & 233.2 & 1 \end{bmatrix}$ | $\begin{bmatrix} 0.82 & -0.57 & -0.02 \\ 0.40 & 0.61 & -0.68 \\ 0.40 & 0.55 & 0.73 \\ -16.98 & -0.07 & 81.37 \end{bmatrix}$ |
| 2 | $\begin{bmatrix} 1133.4 & 0 & 0 \\ 0 & 1138.5 & 0 \\ 345.9 & 244.1 & 1 \end{bmatrix}$ | $\begin{bmatrix} 0.56 & 0.02 & 0.83 \\ 0.62 & -0.67 & -0.4 \\ 0.55 & 0.74 & -0.40 \\ -12.14 & 1.21 & 80.63 \end{bmatrix}$ |
| 3 | $\begin{bmatrix} 1127.7 & 0 & 0 \\ 0 & 1129.7 & 0 \\ 329.2 & 248.6 & 1 \end{bmatrix}$ | $\begin{bmatrix} 0.82 & 0.03 & -0.57 \\ 0.40 & -0.67 & 0.62 \\ 0.40 & -0.74 & 0.54 \\ -17.44 & -11.04 & 87.53 \end{bmatrix}$ |

**SI Figure 1:**
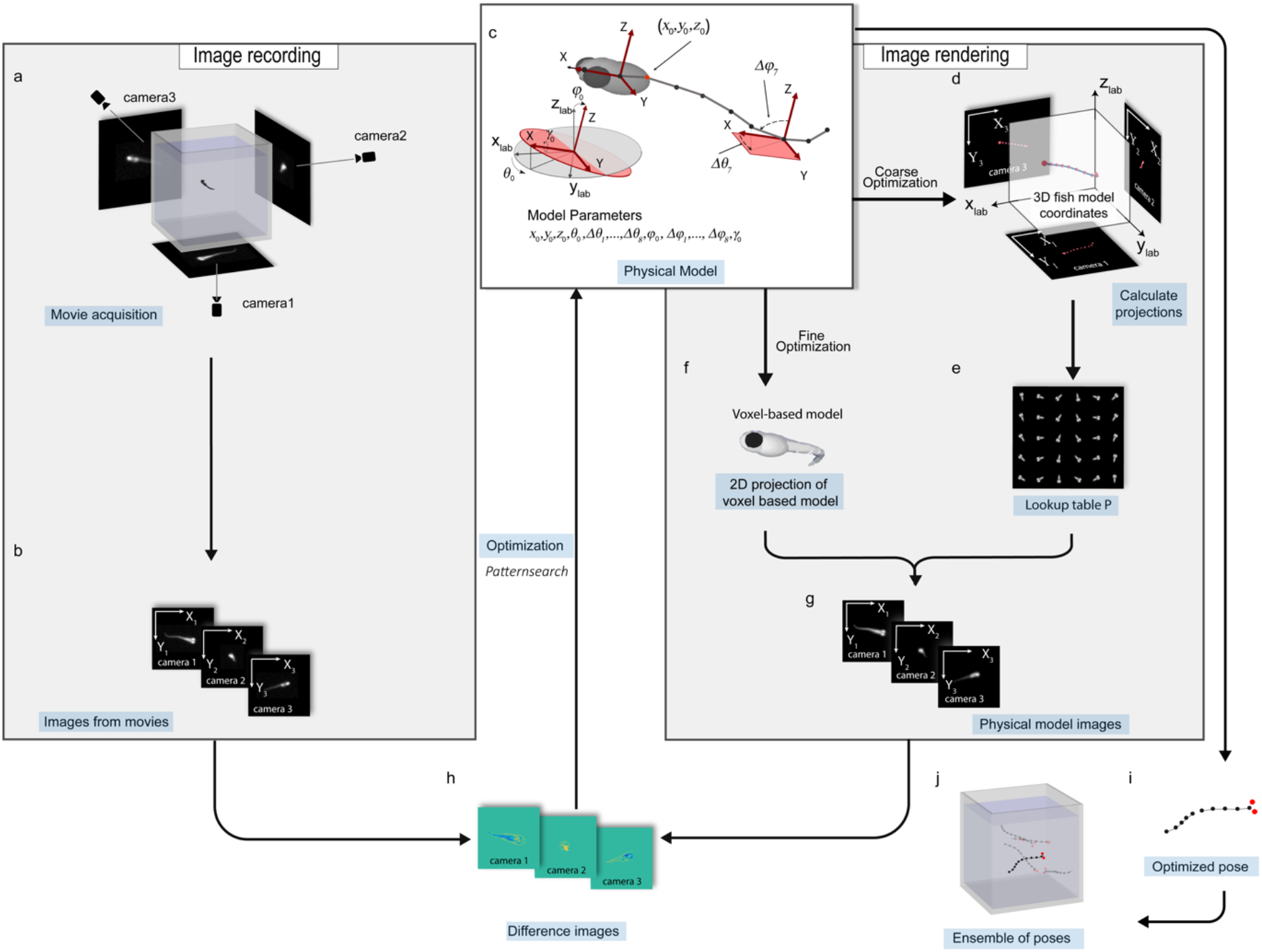
Template-based algorithm tracks 3-D zebrafish swimming poses. (a) Movies of larval swims are acquired at 500 fps by three cameras oriented perpendicular to each other. (b) For each frame, the larval position is triangulated, and the three camera projections are cropped. (c) A model for the larval backbone is generated using 9 rigid segments of equal length oriented at different angles based on an initial guess **p_0_**. Two frames of reference define **p_0_**: (*x*_lab_, *y*_lab_, *z*_lab_) defines the lab reference frame with its origin fixed to the tank’s center and the three axes defined by the principal axes of the cameras. The Euler angles of the larval anterior (*θ*_0_, *φ*_0_, *γ*_0_) are defined with respect to this fixed lab reference frame. (*X, Y, Z*) defines the fish frame of reference fixed to the larval head, with the *X*-axis perpendicular to the line joining the center of the eyes and the *XY*-plane defined by the plane of the eyes. The shape of the larval backbone is parameterized in this reference frame by Δ*θ_i_* and Δ*φ_i_* (see Δ*θ_7_* and Δ*φ_7_* for example). Optimization of the parameter vector **p** occurs in two rounds: *Coarse Optimization* and *Fine Optimization*. (d) Coarse Optimization: The coordinates of the backbone are projected on the three camera planes using their respective calibrated parameters, taking into account refraction. (e) The projections in (d) are used to compute the indices of the appropriate ‘Lookup Table P’ entry rendering a digital projection of each segment of the larva. (f) Fine Optimization: A voxel-based 3-D model is rendered on the three camera planes using the camera calibrated parameters. This step allows a parameter search over **p** at a higher resolution than that achieved using Lookup Table P. (g) The three digital projections are positioned on images with the same resolution as their respective real projections in (b). (h) The sum of the squared pixel values of the three difference images is used as the cost function during optimization. (i-h) The optimization is run until convergence for each frame in parallel to obtain an ensemble of larval poses.

**SI Figure 2:**
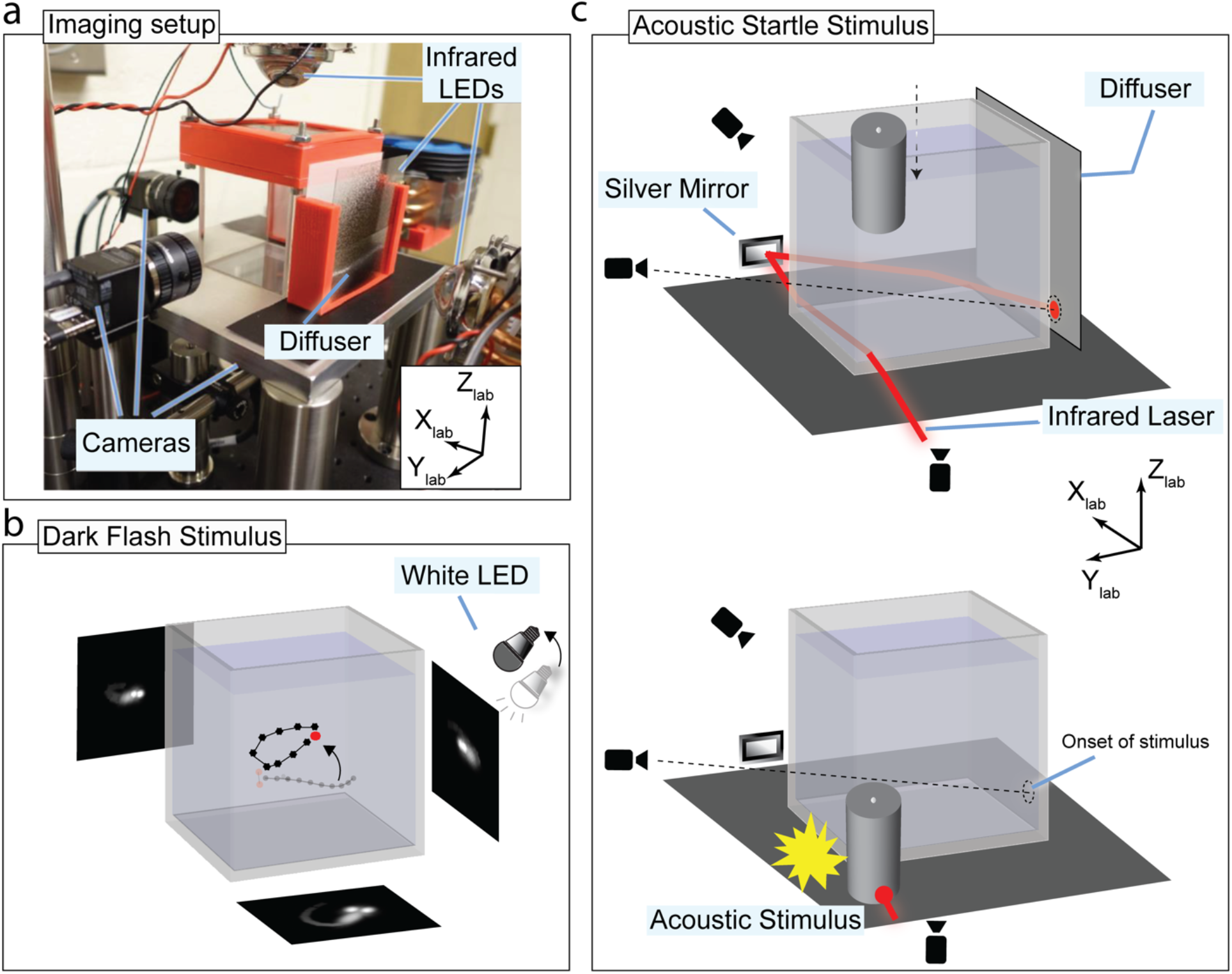
Experimental setup for high-throughput video recording of 3-D larval behavioral experiments. (a) Experimental setup for 3-D experiments. Zebrafish larvae swimming freely in a 7 x 7 x 7 cm glass tank are imaged using three cameras at 500 fps under infrared illumination. (b) Schematic of the setup used to provide acoustic stimulus. Larvae are habituated to white light using a white LED for at least 4 minutes. A dark flash stimulus is provided when the three cameras simultaneously detect a unique larva in their field of view. The instantaneous deactivation of the white LED marks the onset of the dark flash stimulus. (c) Schematic of the setup used to provide acoustic stimulus. The stimulus is generated by dropping a cylindrical weight on the platform supporting the tank from a suitable height. The acoustic signal propagating through the cube startles the larva. The time of the stimulus is recorded using an infrared laser projected in the field of view of one of the cameras. The weight is pulled back up using an Arduino-controlled servo motor and held in place using an electromagnet until the next round of stimulus.

**SI Figure 3:**
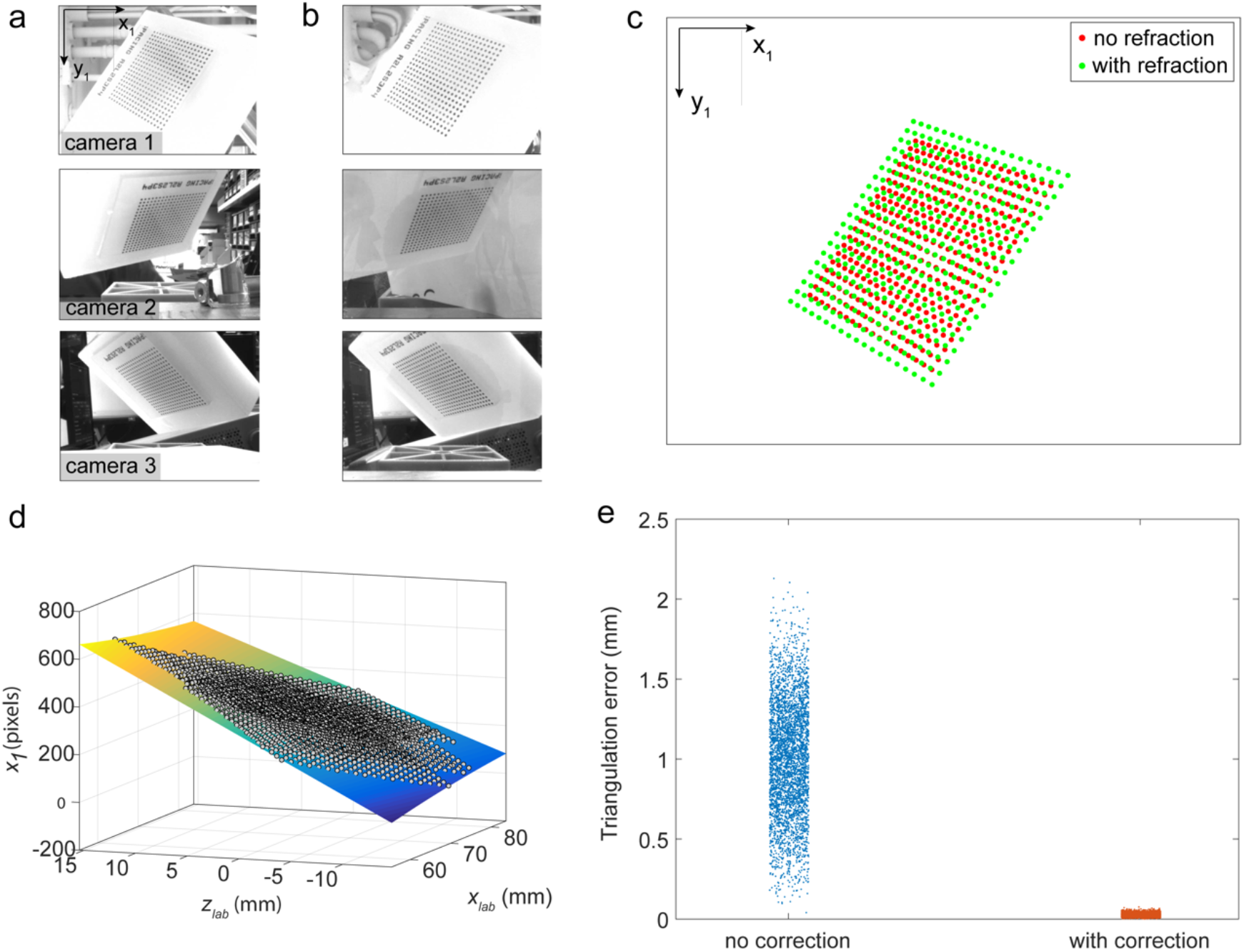
3-D triangulation from 3 camera views with refraction calibration. (a) Images of the calibration dot pattern in air. (b) Images of the same calibration pattern inside the glass cube filled with water. The position of the pattern is identical in both sets of images. (c) A sample image showing the coordinates of the dots on the images taken by one of the cameras. Green dots correspond to an image in (a) (camera 1), in the absence of refraction. Red dots correspond to an image in (b) (camera 1), in the presence of refraction from the glass cube and water. (d) The projection function (*f*_1*x*_ (*y_lab_, z_lab_*)) for the coordinates *x*_1_ in camera 1 of the dot pattern, and fit to a cubic function. The surface plot color represents *x*_1_ for visual convenience. (e) Triangulation error of the lab coordinates of the dots on the calibration pattern from the images taken in water. The mean error is 1.07 mm without refraction correction and 0.03 mm with correction.

**SI Figure 4:**
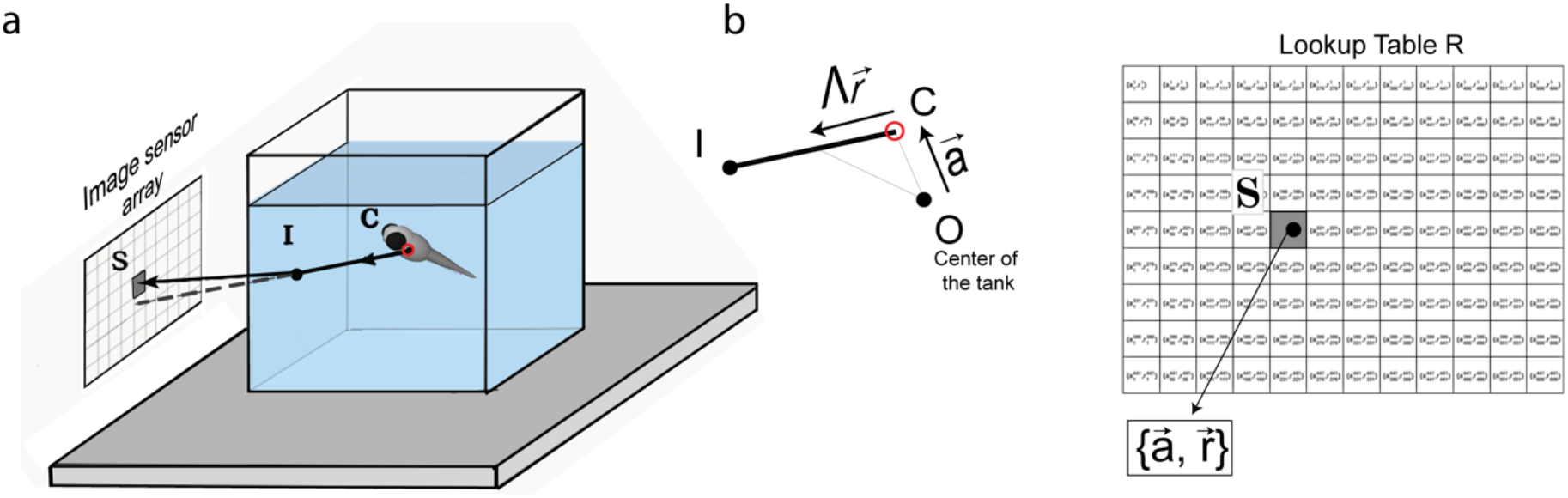
Non-linear projection parameters used to generate Lookup table R. (a) Ray CI emanates from the centroid of the larva, C, and passes through an arbitrary point I in the tank. By invoking ray optics, for every pixel S of the camera sensor array, there is a unique ray CI emanating from a point C inside the tank that is incident on S after refraction at the water-tank-air interface. (b) Lookup Table R stores the two vectors **a** and **r** that uniquely define ray CI for every pixel of the camera sensor, for all three cameras.

**SI Figure 5:**
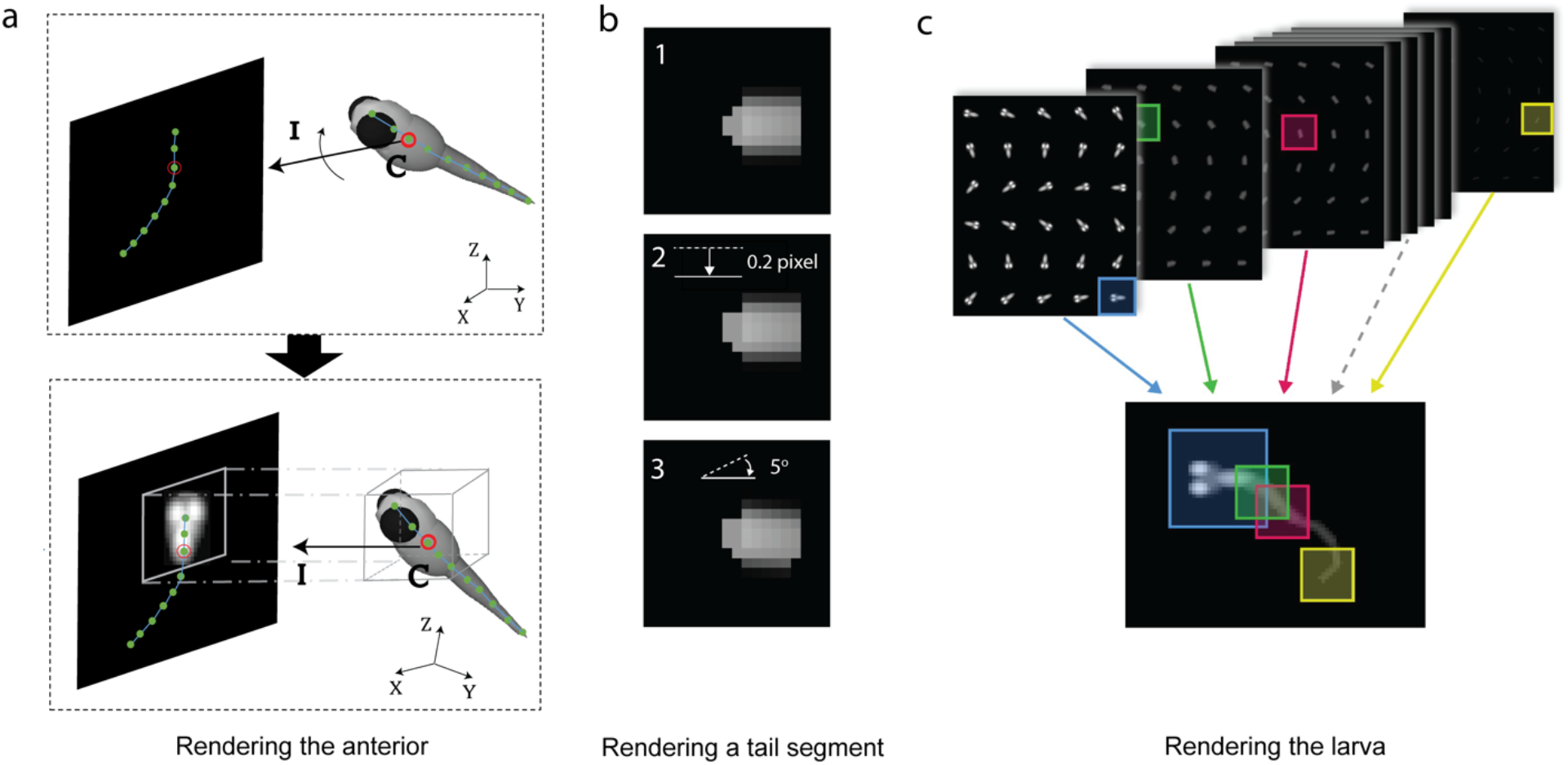
Generation of Lookup Table P entries storing 2-D views. (a) Rendering the larval anterior. Lookup Table P entries of the anterior are generated using orthographic projections of a voxel-based model along the principal axis of each camera. Given the parameter vector **p**, the indices of the appropriate lookup table entry are determined to render the digital projection. The 2-D projection of the larval backbone (green chain) is computed using the nonlinear projection parameters (top panel). Ray CI emanates from the 3-D larva’s centroid and is incident on its 2-D projection. The appropriate lookup table entry to be used is determined by rotating CI such that it is parallel to the principal axis of the corresponding camera (bottom panel). (b) Rendering a tail segment. Image 1 shows a digitally rendered component of a tail segment. Images 2 and 3 show entries in the lookup table adjacent to the entry for image 1. The tail segment in image 2 is 0.2 pixels lower than that in image 1. The segment in image 3 is that in image 1 rotated by 5° clockwise. (c) Rendering the larva. Scheme for the construction of a grayscale fish model from the lookup table. Only a small fraction of the tables is shown.

**SI Figure 6:**
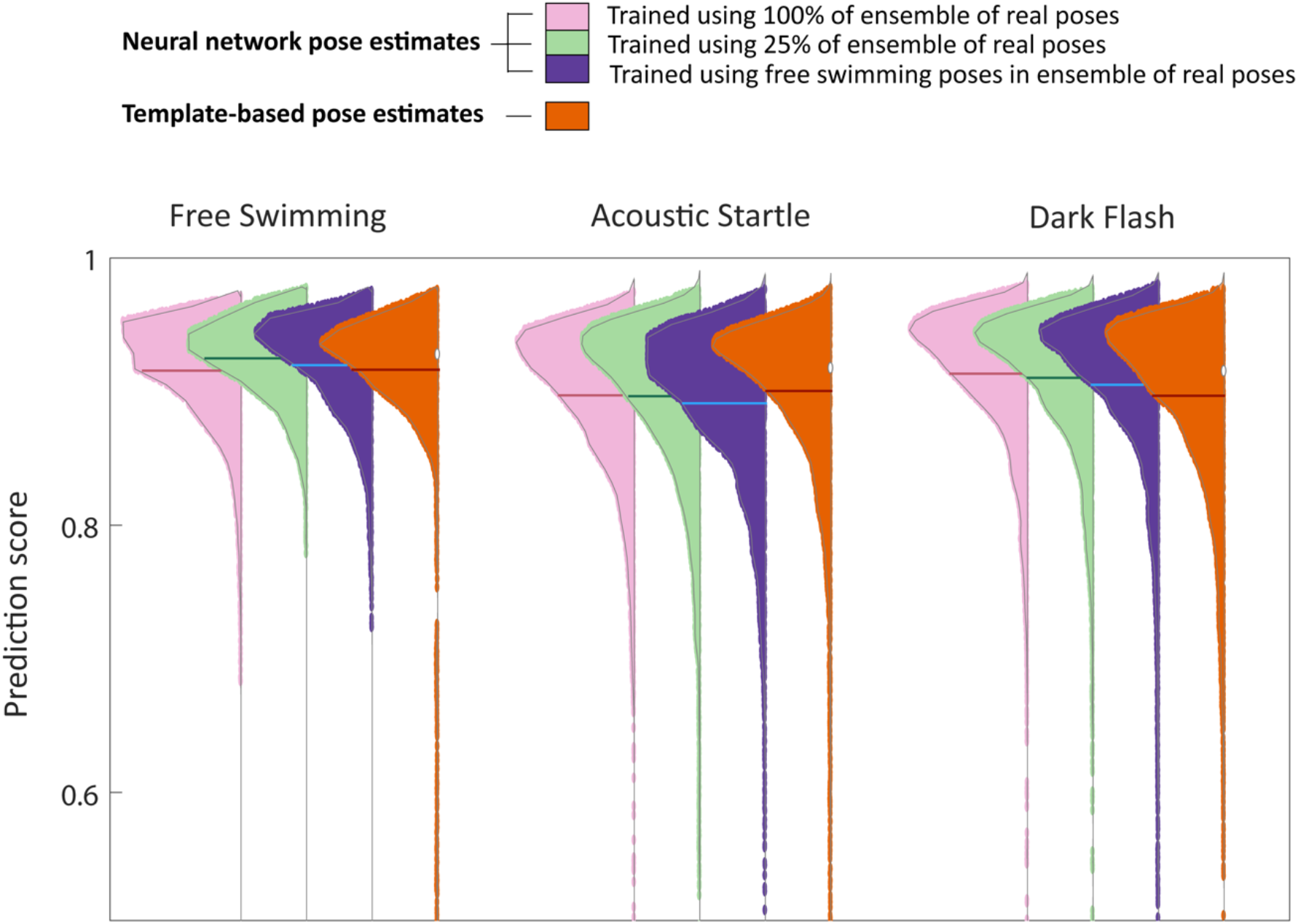
Network predictions are invariant to the amount of training data and comparable to template-based estimates. The performance of the CNN method is not affected by using a smaller subset of real poses to generate the ensemble of physical model poses. The distributions of predictions scores are visualized using kernel density estimate over pose prediction scores obtained using different approaches across free swimming, acoustic startle and dark flash experiments. The distributions compare pose prediction accuracy of network models trained using different subsets of the ensemble of real poses and that of the physical model fits. The mean of each distribution is shown using overlaid dashed lines. Using only 25% (N=8929) of poses from the ensemble of real poses to generate the training dataset does not deteriorate network predictions, compared to the performance of a network trained using 100% (N=35714) of poses from the ensemble of real poses (compare pink and green distributions and their means – overlaid dashed lines). When the training dataset is generated using only free swimming poses from the ensemble of real poses (N=9536), the network performance for acoustic startle and dark flash experiments declines marginally (purple distributions and overlaid dashed lines). The template-based pose estimates are marginally better than neural network predictions in the acoustic startle experiments and consistently worse in dark flash experiments. On average, the network trained using 100% of the data (mean prediction score = 0.9079) is comparable to the template-based predictions (mean predictions score = 0.9029).

